# An InDel Genomic Variant within a Bifunctional Super-Enhancer for LINC00636 and CD47 Regulation in Breast Cancer

**DOI:** 10.1101/2025.11.05.684493

**Authors:** Carolina Di Benedetto, Amelia Tsark, Daniza Acenas, Alysia Thach, Anmol Singhal, Valentina Opazo-Mellado, Anthony Rodriguez Lemus, Mustapha El Zeini, Hui Zhang, Catherine Park, Hugo Gonzalez Velozo, Irving Weissmann, Paola Betancur

**Affiliations:** Department of Radiation Oncology, University of California, San Francisco, San Francisco, CA, USA; Centro Basal Ciencia & Vida, Fundación Ciencia & Vida, Santiago, Chile; Facultad de Ciencias Para el Cuidado de la Salud, Universidad San Sebastian, Santiago, Chile; Department of Anatomy, University of California, San Francisco, San Francisco, CA, USA; Institute for Stem Cell Biology and Regenerative Medicine, Stanford University, Stanford, CA, USA; Ludwig Center for Cancer Stem Cell Research and Medicine, Stanford University, Stanford, CA, USA; Department of Pathology, Stanford University School of Medicine, Stanford, CA, USA

**Keywords:** Super-enhancer, genomic variation, LINC00636, CD47, breast cancer

## Abstract

Highly accessible genomic super-enhancers often drive tumor-promoting programs, yet the impact of genomic variation within these regulatory elements remains unclear. Here, we identified a bifunctional super-enhancer that regulates the expression of cancer-promoting genes LINC00636 and CD47 in breast cancer. We discovered that a common germline insertion variant within the super-enhancer is associated with reduced chromatin accessibility at the super-enhancer locus. Deletion of the insertion in breast cancer cells increased chromatin accessibility, leading to upregulation of LINC00636 and CD47, enhanced resistance to nutrient-deprivation–induced apoptosis (mediated by CD47), activation of senescence (driven by elevated LINC00636), delayed cell death, and reduced infiltration of CD80⁺ pro-inflammatory macrophages, changes that represent tumor-promoting features. Together, our findings uncover a common insertion-deletion variant that fine-tunes the regulatory activity of a bifunctional super-enhancer, suggest a protective role for the insertion allele, and establish a novel function for LINC00636 in senescence and breast cancer.

## INTRODUCTION

Epigenetic modifications that open chromatin and aberrantly expose DNA to transcription factors can dysregulate genes, contributing to cancer development. As a result, these modifications have gained recognition as a hallmark of the disease (1). Super-enhancers (SEs) are large, epigenetically open regions of non-coding DNA, characterized by clusters of cis-regulatory elements densely occupied by transcription factors and coactivators, as well as active histone marks (2,3). Initially identified as key regulators of gene expression programs critical for cell identity, SEs have since been shown to also control the expression of oncogenes and other genes that confer growth advantages to tumor cells (2,4,5). We have previously shown that SEs are linked to genes involved in chemotherapy resistance in breast cancer (6), as well as to anti-apoptotic genes in both breast and lung cancer cells (7).

Given their central role in regulating cancer-promoting genes, SEs have been considered promising therapeutic targets, leading to the development of small molecule inhibitors, such as Bromodomain and Extra-Terminal (BET) inhibitors, which disrupt the binding of bromodomain-containing proteins to acetylated histones and suppress the expression of SE-regulated genes (3). These inhibitors showed promising efficacy in preclinical models of various cancers, including breast cancer (3,8,9). However, therapeutic targeting of SEs remains challenging because of their large genomic size and their essential roles in regulating gene expression in normal cells. Therefore, it is crucial to identify alternative strategies for specifically targeting SEs that regulate cancer-promoting genes. Recent studies have identified core elements—smaller, specific DNA regions—within active SEs that are essential for SE function in distinct cell types (10–13). These findings provide valuable insights for developing targeted approaches to disrupt SE-driven gene overexpression and address transcriptional dysregulation, specifically in cancer cells.

While our understanding of how cis-regulatory elements—particularly those within SEs— modulate gene expression has advanced significantly, the role of genomic variation, including germline variants within regulatory enhancers, in tumorigenesis remains poorly understood. Genetic variation, such as single nucleotide polymorphisms (SNPs) and insertions or deletions of small DNA fragments (InDels), within core elements of SEs can lead to changes in gene expression patterns contributing to cancer susceptibility or protection. For instance, a common SNP that reduces GATA3 binding to the SE region of the proliferation-promoting LMO1 gene in pediatric malignancies, decreases LMO1 levels and acts as a protective SNP (14,15).

Understanding how genetic variation within SEs regulates cancer-promoting genes is key to uncovering the mechanisms that drive either cancer progression or protection. This knowledge could ultimately guide the development of targeted anti-cancer therapies tailored to individual patients based on their variant status.

Here, we report that a SE region located between LINC00636, a long intergenic non-coding RNA, and CD47, a cell surface receptor that signals ‘don’t eat me’ to macrophages, upregulates not only CD47 (13) but also neighboring LINC00636 in breast cancer. Within this bifunctional SE, we identified a common germline InDel variant, where the insertion allele is linked to reduced chromatin accessibility at the SE locus, based on analyses of genomic data from breast cancer patients available in The Cancer Genome Atlas (TCGA). By performing Assay for Transposase-Accessible Chromatin with high-throughput sequencing (ATAC-seq) on an engineered breast cancer cell line in which we deleted the insertion, we confirmed that removal of the insertion increases SE chromatin accessibility and subsequently upregulates LINC00636 and CD47 expression. Furthermore, deletion of the insertion promoted a tumorigenic phenotype both in vitro and in vivo. We observed increased cell death in spheroid models carrying the insertion and apoptosis upon glucose deprivation. By contrast, deletion of the insertion led to a CD47-mediated reduction in apoptosis, whereas overexpression of LINC00636 activated a senescence program in breast tumor cells. In xenograft models, tumors harboring the insertion exhibited greater infiltration of pro-inflammatory macrophages, while absence of the insertion reduced the proportion of these macrophages—a change associated with enhanced cancer cell survival and tumor progression. Finally, transcriptomic analysis revealed a downstream pro-tumoral gene program—encompassing apoptosis resistance, therapy resistance, senescence, and immune evasion—triggered by deletion of the insertion variant and the subsequent upregulation of LINC00636 in breast cancer cells. Our findings demonstrate that a common InDel variant modulates the bifunctional activity of a SE regulatory region. In the absence of the insertion, LINC00636 and CD47 are upregulated, driving pro-tumoral changes and enhancing immune evasion. In contrast, the insertion acts protectively by downregulating both genes. For the first time, we highlight the differential function of this InDel variant, underlining its potential as a protective biomarker for attenuating a SE-driven program of tumor cell survival. In addition, our study reveals a novel role for LINC00636 in breast cancer.

## RESULTS

### LINC00636 and CD47 gene regulation is associated with a patient-specific highly accessible intergenic regulatory region in breast tumors

Using Chromatin Immunoprecipitation followed by high-throughput sequencing (ChIP-seq) with antibodies against H3K27ac––a mark of active enhancers (16)––and functional assays, we previously identified a SE regulatory region. This region is located between the CD47 gene and the long intergenic non-coding RNA LINC00636 in breast cancer cell lines and a xenograft model (13). To investigate whether this large regulatory region is specific to individual patients, we analyzed ATAC-seq data from 74 patient-derived breast tumor samples available in the TCGA database. We employed the Rank Ordering of Super-Enhancers (ROSE) algorithm, which combines nearby accessible DNA elements into clusters. These clusters are then ranked according to their accessibility signal density in each tumor sample. Next, we applied the method developed by White et al (17) to distinguish the most highly accessible regions in each breast tumor (Figure 1A). Surprisingly, we found that the intergenic regulatory region between LINC00636 and CD47 is highly accessible (Figure 1B) in 50% of the breast tumors analyzed. Moreover, using the TCGA breast cancer ATAC-seq dataset and available corresponding gene expression data, we found that there is a positive and significant correlation between chromatin accessibility of the intergenic regulatory region and LINC00636 and CD47 RNA expression levels in breast tumors (Figure 1C). These findings suggest that chromatin opening and increase in accessibility of this intergenic region upregulate the expression of LINC00636 and CD47 in breast cancer. In addition, our analyses suggest that, for a large percentage of breast cancer patients, a highly accessible intergenic regulatory region may be associated with LINC00636 and CD47 gene upregulation. It is well known that more accessible chromatin at enhancers facilitates efficient binding of transcription factors, which recruit additional proteins to bridge enhancers and promoters, thereby enhancing transcription (2). In line with this, we found that the intergenic regulatory region between LINC00636 and CD47 physically interacts with the promoter region of LINC00636 (Figure 1D). To discern this interaction, we mined the GeneHancer database of human enhancers, which links enhancers to genes using multi-source genomic data including Capture Hi-C promoter-enhancer long range interactions (18). This analysis suggests a direct regulatory role for this intergenic regulatory region on LINC00636 expression not previously reported.

**Figure 1.**
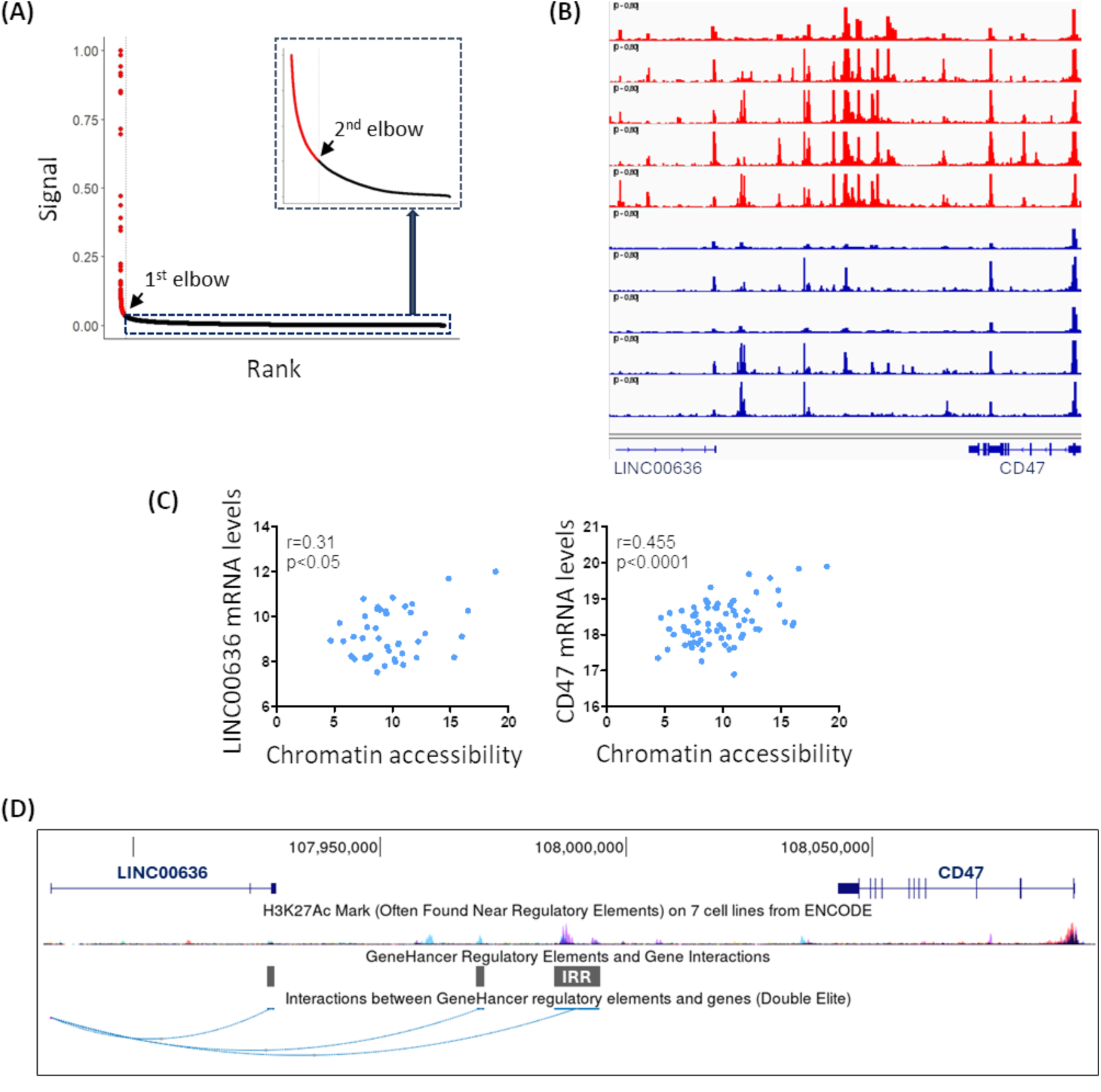
LINC00636 and CD47 gene regulation is associated with a patient-specific highly accessible intergenic regulatory region in breast tumors. (A) Representative ranking of ATAC-seq accessible regions in one breast cancer patient following White’s method. All accessible regions within a 12.5kb distance were combined into clustered regions and clustered regions were ranked based on their accessibility using ROSE. To identify highly accessible regions, the accessibility signal for each cluster was plotted against the clusters’ rank, and the elbow point (first inflection point) was calculated. Next, we calculated a second elbow point for the lower portion of the curve (highlighted in a blue dashed rectangle), using the first inflection point as the new maximum value. Clusters whose signal was higher than the second elbow point were considered highly accessible (shown in red). (B) IGV view of ATAC-seq enrichment data of the intergenic region between LINC00636 and CD47 for TCGA breast tumors. The 5 samples with most accessible or least accessible DNA at the intergenic regulatory region are shown. (C) LINC00636 and CD47 RNA levels significantly correlate with chromatin accessibility of the intergenic regulatory region in TCGA breast tumors. n=45 for LINC00636 and n=72 for CD47. Pearson correlation analysis was performed. r=Pearson correlation coefficient. (D) Publicly available GeneHancer data show that the intergenic regulatory region (IRR) physically interacts with LINC00636 TSS. Data was obtained and visualized on UCSC genome browser. Hg38 was used as reference genome. Ruler shows location on chromosome 3.

### LINC00636 and CD47 gene expression is regulated by a bifunctional SE

We previously showed that the highly accessible regulatory region between LINC00636 and CD47 is a SE in the breast cancer cell lines MCF7 and HCC1954 (13). To experimentally demonstrate that LINC00636 is upregulated by large regulatory regions such as SEs, we disrupted its regulatory function by treating breast cancer cell lines carrying or lacking the SE with BET inhibitors and then analyzed LINC00636 RNA expression after 6 and 24 hours. BRD4, a member of the BET protein family, works as a bridge between hyperacetylated chromatin regions, such as SEs, and the TEFb (transcription elongation factor) complex, therefore, enhancing transcription (8). We observed that treatment with two different BRD4 inhibitors, JQ1 (19) and I-BET151 (20), reduces LINC00636 RNA levels to a greater extent than CD47, particularly in the MCF7 and HCC1954 breast cancer cell lines, which carry the SE. This effect was more pronounced compared to breast cancer cell lines BT549 and T47D that lack the SE (Figure 2A and S1), confirming that LINC00636 gene expression is regulated by SEs in breast cancer cells. Next, to determine experimentally whether the SE region interacts with LINC00636 and regulates its expression, we employed a CRISPR activation (CRISPRa) system (Figure 2B). This system uses a deactivated Cas9 (dCas9) fused to the VP64 transcriptional activator domain (a tetramer of VP16) (21,22), and a guide RNA that directs dCas9-VP64 to the target site (at chr3:107987579-107987598), corresponding to the previously identified active core element E5 within the SE (13). If the SE is functional, activation through E5 should increase transcription of LINC00636—and, as previously shown, CD47—via enhancer–promoter looping that brings VP64 into proximity with the transcriptional start sites (TSSs) of both genes (23). Indeed, transiently targeting dCas9-VP64 to the SE significantly increased LINC00636 and CD47 RNA expression when compared to targeting a random region (Figure 2B). We noticed that this effect is greater in breast cancer cells MCF7 and HCC1954, which carry the SE, when compared to those that lack the SE. Consistent with our previous observations, we also noticed that the change in gene expression levels upon SE-activation in breast cancer cells was higher for LINC00636 than for CD47.

**Figure 2.**
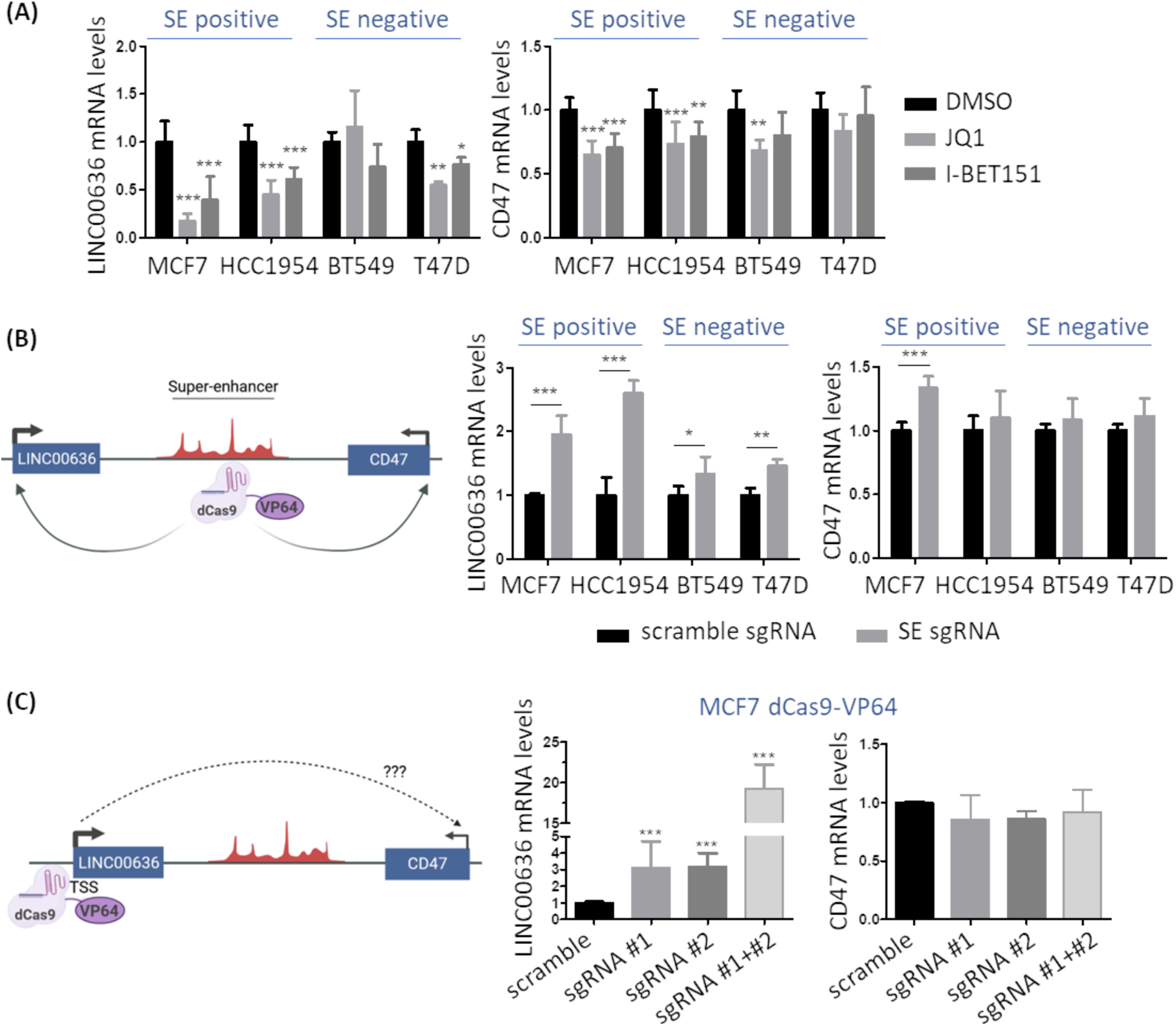
LINC00636 RNA expression is regulated by a bifunctional SE. (A) LINC00636 and CD47 RNA expression levels decrease upon BET inhibition in BCCL, and this reduction occurs at a greater extent in the cells where the SE is present. Cells were treated with 1 μM JQ1 or I-BET151 for 6 hours and mRNA levels were analyzed by qPCR. DMSO was used as vehicle control. Two-way ANOVA was performed, *p<0.05, **p<0.01, ***p<0.001. (B) CRISPRa of the SE locus increases LINC00636 and CD47 RNA expression levels, and this activation occurs at a greater extent in the cells where the SE is present. dCas9-VP64-expressing cells were transfected with scramble sgRNA or sgRNA targeting the E5 core element within the SE. mRNA levels were analyzed by qPCR 48 hours after transfection. Two-way ANOVA was performed, *p<0.05, **p<0.01, ***p<0.001. (C) LINC00636 overexpression does not affect CD47 RNA expression levels. Left image shows the CRISPRa approach used for LINC00636 overexpression targeting LINC00636 TSS. The plots show LINC00636 and CD47 RNA levels in dCas9-VP64-expressing MCF7 cells transfected with sgRNAs targeting two different sites on LINC00636 TSS. mRNA levels were analyzed by qPCR 48 hours after transfection. ANOVA was performed, *p<0.05, **p<0.01, ***p<0.001.

Long intergenic non-coding RNAs can regulate gene expression by both cis and trans mechanisms (24). Since we noticed a stronger regulatory effect on LINC00636 when targeting the SE with CRISPRa, we speculated that SE-driven CD47 regulation could be indirect and through modulation of LINC00636 RNA levels. To test this, we transiently overexpressed LINC00636 in MCF7 cells using CRISPRa along with sgRNAs that target LINC00636 TSS. This approach has the advantage of increasing LINC00636 expression endogenously, which is crucial to examine its cis-acting effects. LINC00636 overexpression through this approach did not affect CD47 RNA levels in MCF7 cells (Figure 2C), thus indicating that CD47-regulation by the SE is not mediated by LINC00636. Overall, we show that LINC00636 and CD47 are regulated by a bifunctional SE in breast cancer cells.

### An Insertion/Deletion (InDel) germline variant linked to chromatin accessibility is located within the bifunctional SE

Core elements within SEs have recently been described as key players of SE-mediated gene regulation (10–13). To investigate whether core elements within the intergenic region between LINC00636 and CD47 drive their expression in breast cancer, we correlated ATAC-seq chromatin accessibility data of all accessible regions in the LINC00636-CD47 neighborhood identified on TCGA breast tumors by Corces et al (25) with matching RNA sequencing (RNA-seq) gene expression for each patient. We observed that, whereas CD47 gene expression positively and significantly correlates with chromatin accessibility at most accessible regions across the region analyzed, LINC00636 expression seems to be more dependent on the activity of fewer regulatory elements (Figure 3A). Moreover, within the bifunctional SE region, only accessibility of peak 10 significantly correlates with both LINC00636 and CD47 RNA levels. Notably, peak 10 overlaps with a core element we previously named E5, which we experimentally demonstrated is critical for SE-driven CD47 upregulation in certain breast cancers (13). Our new computational findings indicate that E5 plays a key role in enhancing the expression of both genes. Next, we performed targeted PCR sequencing at the E5 element in breast cancer samples and, within it, found a biallelic 8-bp insertion/deletion variant (InDel) (Figure 3B). Analysis of genomic datasets from 75,699 individuals showed that the InDel is a common germline variant (rs58296163, located on chr3, 3-107987596-TTTGGGAC-(GRCh38)), with a minor allele frequency (MAF) of 46.5% for the insertion, and a major allele frequency of 53.5% for the deletion (Figure 3C). We noticed that allele frequencies varied substantially across populations. For instance, in non-Finnish Europeans, the insertion is the major allele (53.7%) while the deletion is the minor allele (46.3%). In contrast, the deletion is the major allele in other populations—for example, in East Asians (88% deletion, 12% insertion) and in Africans/African Americans (61.8% deletion, 38.2% insertion). Previous reports have shown that genomic variation can modulate the chromatin landscape or SEs activity, thereby affecting their regulatory function, as demonstrated by a protective SNP in the LMO1 oncogene’s SE that reduces chromatin accessibility and GATA3 binding, leading to lower LMO1 expression and decreased cell proliferation (14,15). To investigate whether the InDel variant correlates with the accessibility of the bifunctional SE, we first classified a cohort of TCGA breast tumors as homozygous for either the insertion or the deletion, based on DNA sequences obtained from ATAC-seq reads covering the InDel region. Then, we analyzed ATAC-seq enrichment profiles in the same TCGA breast cancer patients. We observed that enhancer chromatin within the SE locus is significantly less accessible in breast cancer patients that are homozygous for the insertion allele compared to patients that are homozygous for the deletion (Figure 3D). Thus, we identified an InDel germline variant linked to bifunctional SE accessibility in breast cancer patients.

**Figure 3.**
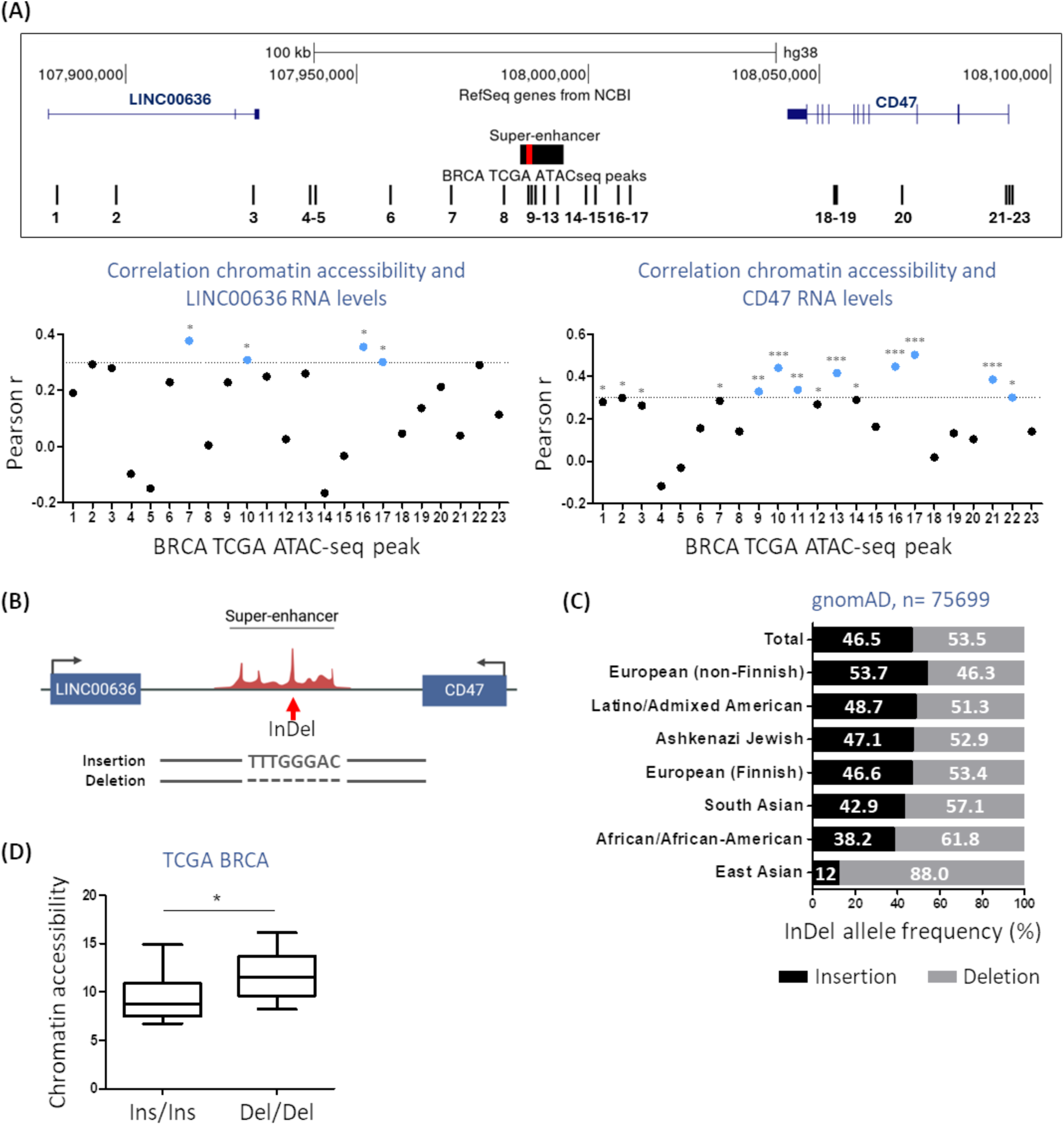
An InDel variant is located within the bifunctional SE. (A) Chromatin accessibility of specific enhancers highly correlate with LINC00636 and CD47 RNA expression in TCGA breast tumors. Top panel shows location of the bifunctional SE and of all TCGA ATAC-seq peaks in the LINC00636-CD47 neighborhood. The E5 core element within the SE is marked in red. Peaks 9 to 13 are located within the bifunctional SE. Graphs represent the Pearson correlation coefficient, r, between chromatin accessibility and LINC00636 or CD47 RNA levels for each peak. Peaks with r>0.3 are highlighted in blue. n=45 for LINC00636 and n=72 for CD47. (B) SE genomic locus showing the nucleotide sequence of the insertion and deletion alleles. (C) InDel allele frequency across populations, grouped by ethnicity. Publicly available data for n=75699 obtained from gnomAD, n=33838 European (non-Finnish), n=7612 Latino/Admixed American, n=1730 Ashkenazi Jewish, n=5271 European (Finnish), n=2405 South Asian, n=20638 African/African American, n=2558 East Asian. Other under-represented ethnicities are not displayed (n=1647). (D) Chromatin at the SE locus is less accessible in breast cancer patients that are homozygous for the insertion compared to those homozygous for the deletion. TCGA BRCA, n=27. Two-tailed unpaired t-test was performed, *p<0.05.

### Deletion of the insertion increases chromatin accessibility at the SE locus and drives LINC00636 and CD47 overexpression in breast cancer cells

To experimentally test if deleting the insertion increases chromatin accessibility at the SE locus and hence increases expression of LINC00636 and CD47 target genes, we used CRISPR editing to remove the insertion in MCF7 breast cancer cells. MCF7 cells are a suitable model for these studies since they harbor the SE and are homozygous for the insertion. We successfully isolated two clones, MCF7Δ#1 and MCF7Δ#2, in which the insertion was completely deleted in the two alleles (Figure 4A). We performed ATAC-seq in the parental MCF7 cells carrying the insertion or MCF7Δ#1 and MCF7Δ#2 clones carrying the engineered deletion. Our data shows that chromatin at the SE locus is more accessible in the MCF7Δ#1 and MCF7Δ#2 clones carrying the engineered deletion, i.e., chromatin at the SE locus is less accessible when the insertion is present, and this effect in accessibility is specific to the SE region since we did not observe a similar effect near promoter regions (Figure 4B). In line with these findings, LINC00636 and CD47 expression levels were higher in the MCF7Δ#1 and MCF7Δ#2 clones compared to MCF7 parental cells, in which we observed reduced expression levels of both genes (Figure 4C). In a different luminal breast cancer cell line T47D, deletion of the insertion increased CD47 expression but did not affect LINC00636 (Figure S2). By contrast, deletion of the insertion in HCC1954 cells (HER2+, InDel heterozygous) or in BT20 and MDAMB231 cells (both basal, homozygous for the insertion) had no effect on either gene (Figure S2). These findings across different breast cancer cell lines suggest that SE accessibility and subtype-specific regulatory factors influence how the InDel variant modulates LINC00636 and CD47. Taken together, our results indicate that loss of the insertion increases chromatin accessibility of the bifunctional SE, whereas its presence reduces accessibility—likely by recruiting repressive factors that close chromatin at the SE locus and thereby attenuate expression of SE-associated genes LINC00636 and CD47 (Figure 4D).

**Figure 4.**
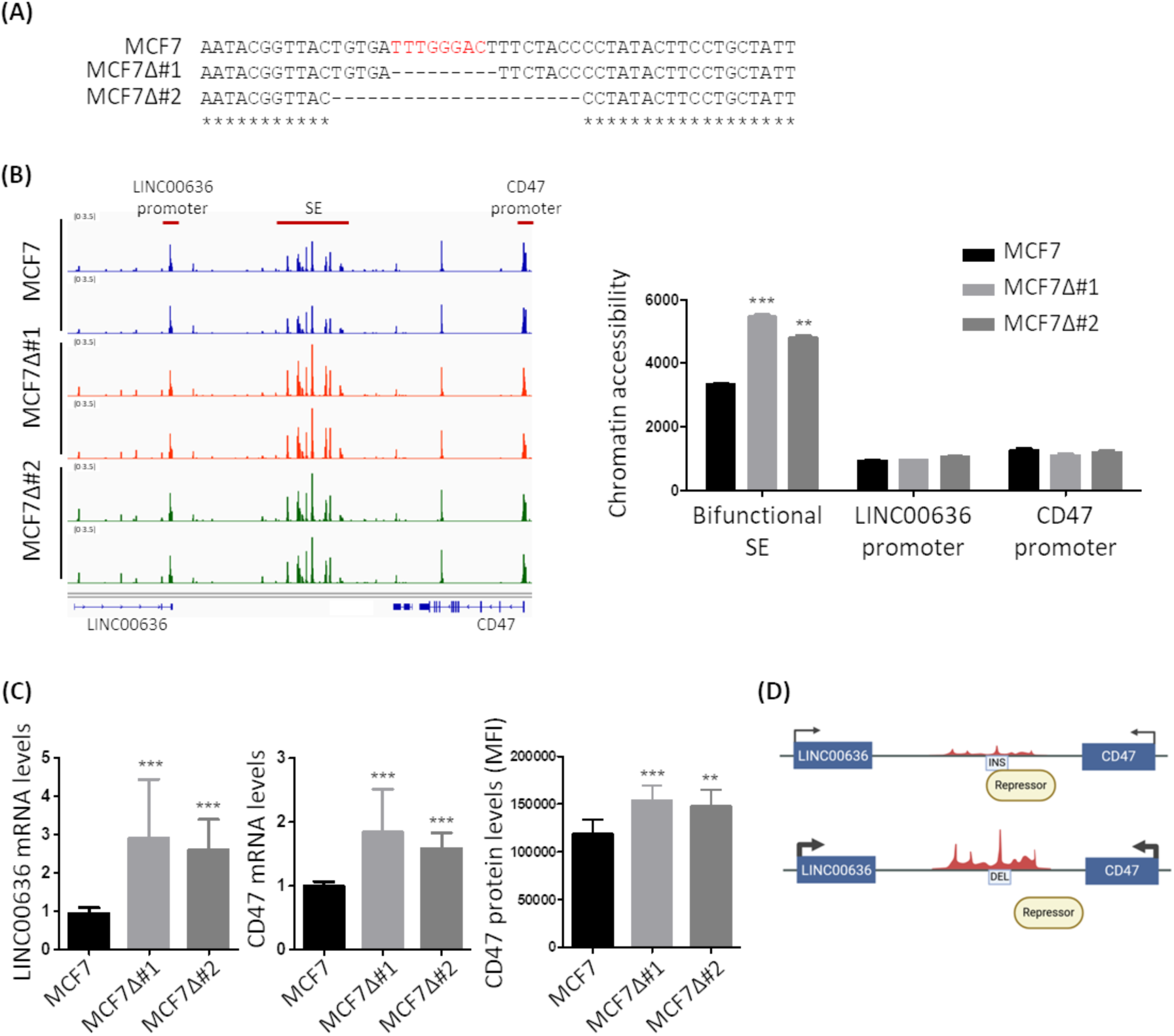
Deletion of the insertion increases chromatin accessibility at the SE locus and drives LINC00636 and CD47 overexpression in MCF7 cells. (A) DNA alignment at the insertion locus in MCF7 and the MCF7Δ#1 and MCF7Δ#2 clones isolated upon deleting the insertion variant by CRISPR. Alignment of DNA sequences was performed using CLUSTAL O (1.2.4). The insertion is highlighted in red. Nucleotides that are conserved across the three samples are marked with an asterisk. (B) Chromatin at the SE locus is more accessible in MCF7Δ#1 and MCF7Δ#2 clones compared to MCF7 parental cells. On the contrary, chromatin accessibility at LINC00636 and CD47 promoters is not affected by deleting the insertion variant. ATAC-seq experiments were performed. ANOVA was performed, **p<0.01, ***p<0.001, n=2. (C) LINC00636 RNA levels and CD47 RNA and protein levels are higher in MCF7Δ#1 and MCF7Δ#2 clones compared to MCF7 parental cells. ANOVA was performed, **p<0.01, ***p<0.001. (D) Model showing that the insertion allele is a binding site for a repressive factor. When the insertion is absent, the repressive factor does not bind to the InDel locus, the SE region becomes more accessible and LINC00636 and CD47 gene transcription increases.

### Deletion of the insertion enhances resistance to cellular stress and reduces infiltration of pro-inflammatory macrophages

To investigate the functional role of the insertion variant in breast cancer phenotypes, we generated 3D tumor spheroid models using parental MCF7 cells carrying the insertion or MCF7Δ#1 and MCF7Δ#2 clones carrying the engineered deletion. 3D tumor spheroids offer a physiologically relevant in vitro model for studying tumor behavior, as they recapitulate key features of the tumor microenvironment, including cell-cell interactions, hypoxia, and gradients of nutrient availability (26). We monitored 3D spheroid formation and growth over time and observed a distinct morphological difference that resembled an increase in cell death in spheroids generated from the parental MCF7-derived spheroids compared to MCF7Δ#1 or MCF7Δ#2 clones. While the spheroids generated from the parental line were characterized by the release of individual cells and the loss of a tightly compact, spherical structure beginning on Day 7 (Figure 5A), the ones generated from MCF7Δ#1 and MCF7Δ#2 cells maintained their spheroid integrity at this timepoint with no visible signs of cell death. The cell death phenotype observed in the parental MCF7 spheroids emerged later in the MCF7Δ#1 and MCF7Δ#2 spheroids on Day 9 (Figure S3A). To confirm that the phenotype was associated with delayed cell death, we stained the spheroids with BD Via-Probe™ Green, a nucleic acid stain specific for dead cells. Fluorescent microscopy showed significantly fewer dead cells in MCF7Δ#1 and MCF7Δ#2 spheroids compared to parental MCF7 spheroids (Figure 5B). In addition, 3D cultures derived from MCF7 cells with stable CRISPRa activation of the bifunctional SE—achieved by targeting the E5 core element (chr3:107987579–107987598; Figure 2B) adjacent to the region harboring the InDel—maintained spheroid integrity relative to control cells (Figure 5C). This phenotype resembled that observed in the MCF7Δ#1 and MCF7Δ#2 clones lacking the insertion. Overall, our CRISPRa experiments confirm that the E5 element, which harbors the InDel variant, contributes to the spheroid phenotype observed in the absence of the insertion. Because stable CRISPRa activation of this element also increases LINC00636 and CD47 expression (Figure S3B), elevated levels of these genes are likely mediating the phenotype.

**Figure 5.**
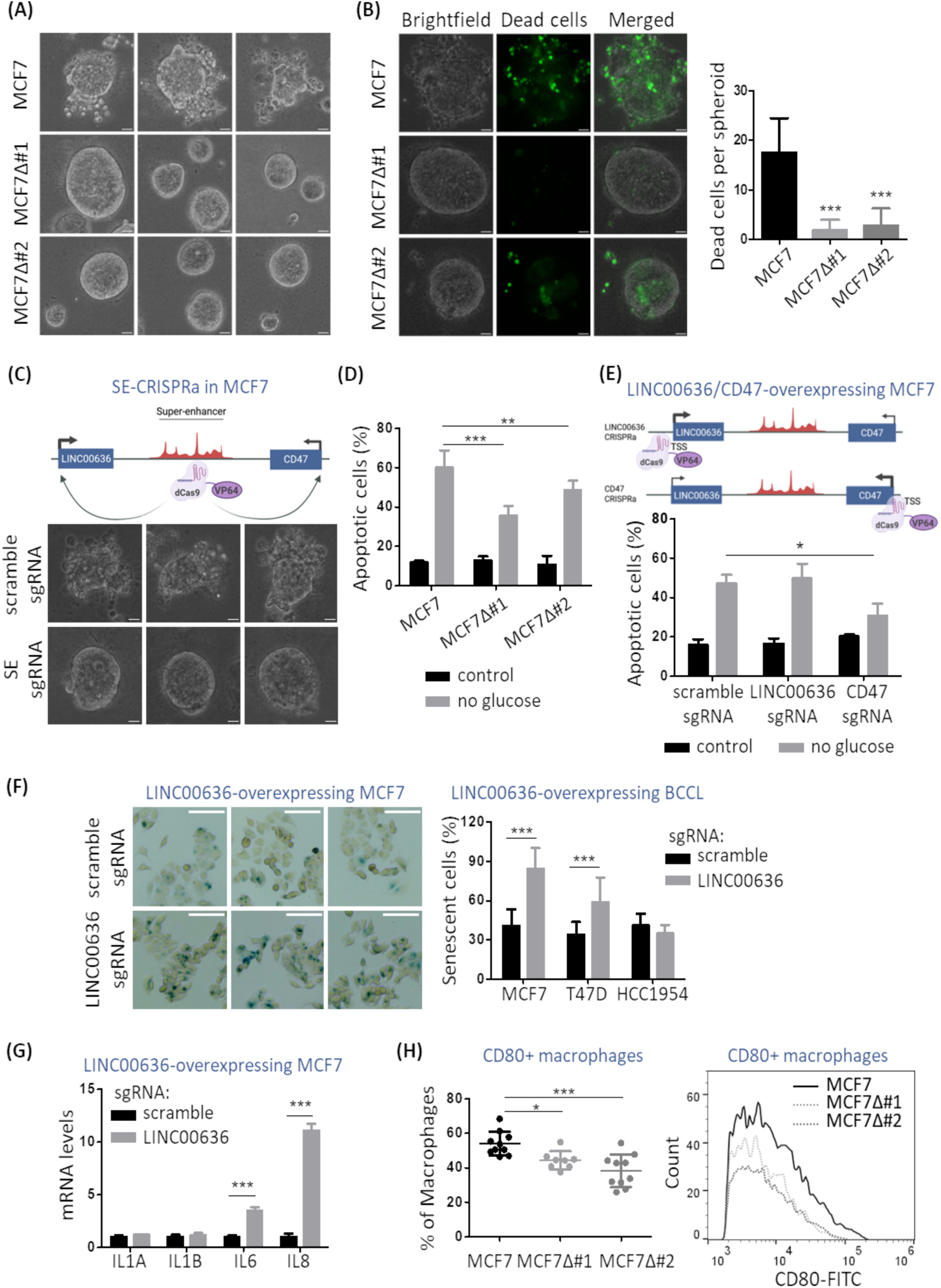
Deletion of the insertion enhances resistance to cellular stress and reduces infiltration of pro-inflammatory macrophages. (A) MCF7Δ#1 and MCF7Δ#2 clones maintain spheroid integrity in 3D cultures, while MCF7 cells lose compact spherical morphology. MCF7Δ#1 and MCF7Δ#2 clones or MCF7 control cells were seeded on plates pre-coated with Matrigel and imaged at day 7. Three representative images per condition are shown. Scale bars represent 100 μm. (B) Deleting the insertion reduces cell death in 3D cultures. MCF7Δ#1 and MCF7Δ#2 clones or MCF7 control cells were seeded on plates pre-coated with Matrigel and imaged at day 7 upon staining with BD Via-Probe™ Green Nucleic Acid Stain. Panels show bright field, fluorescence (cell death) and merged images. Scale bars represent 100 μm. Number of dead cells per spheroid was quantified using ImageJ. (C) CRISPRa of the bifunctional SE in MCF7 cells restores spheroid integrity in 3D cultures. MCF7 cells expressing dCas9-VP64 and sgRNAs scramble control or targeting the E5 core element within the SE were seeded on plates pre-coated with Matrigel and imaged on day 7. Top image shows the CRISPRa strategy used to activate the bifunctional SE. Scale bars represent 100 μm. (D) Deleting the insertion increases resistance to apoptosis induced by glucose deprivation in MCF7. MCF7Δ#1 and MCF7Δ#2 clones or MCF7 control cells were grown in media without glucose for 48 hours and the percentage of apoptotic cells was quantified by Annexin V staining and flow cytometry. Two-way ANOVA, **p<0.01, ***p<0.001. (E) CD47 overexpression increases resistance to apoptosis induced by glucose deprivation in MCF7. MCF7 cells expressing dCas9-VP64 and sgRNAs scramble control or sRNAs targeting LINC00636 or CD47 promoters were grown in media without glucose for 48 hours and the percentage of apoptotic cells was quantified by Annexin V staining and flow cytometry. Top image shows the CRISPRa strategy used for LINC00636 or CD47 overexpression. Two-way ANOVA, *p<0.05. (F) LINC00636 overexpression increases senescence in breast cancer cells. MCF7, T47D or HCC1954 cells expressing dCas9-VP64 and sgRNAs scramble control or sgRNAs targeting LINC00636 TSS were studied. 48 hours post-seeding, senescence was analyzed by b-galactosidase staining, images were captured, and the percentage of senescent (blue) cells was quantified. Left panel shows representative images of MCF7 cells overexpressing LINC00636 or control. Scale bars represent 110 μm. Two-way ANOVA, ***p<0.001. (G) LINC00636 overexpression increases expression of senescence markers in MCF7. MCF7 cells expressing dCas9-VP64 and sgRNAs scramble control or sgRNAs targeting LINC00636 TSS were studied. 48 hours post-seeding, senescence markers IL1A, IL1B, IL6 and IL8 were analyzed by qPCR. Two-way ANOVA, ***p<0.001. (H) Infiltration of CD80+ macrophages decreases in tumors derived from insertion-deleted MCF7Δ#1 and MCF7Δ#2 clones compared to parental MCF7 cells. CD80 expression was analyzed by flow cytometry upon collection of the breast tumors. Left graph shows CD80+ cells infiltrating the tumors as percentage of macrophages. Right graph is the histogram of CD80-FITC fluorescence in CD80+ macrophages. ANOVA, *p<0.05, ***p<0.001.

To test whether the integral spheroid morphology observed in MCF7Δ#1 and MCF7Δ#2 clones, as well as upon CRISPRa activation of the InDel-containing region within the bifunctional SE, is driven by high levels of LINC00636 or CD47, we next stably overexpressed LINC00636 or CD47 in MCF7 cells using CRISPRa with sgRNAs targeting their respective TSSs (or a scramble sgRNA as control). As shown in Figure S3C, overexpression of either gene alone was not sufficient to maintain spheroid integrity on day 7.

Given that nutrient limitation triggers cell death in 3D cultures, we next asked whether the increased expression of LINC00636 or CD47 contributes to the integral spheroid morphology and cell death delay observed in MCF7Δ#1 and MCF7Δ#2 clones by providing resistance to nutrient-deprivation–induced cell death. To test this, we first assessed whether the deletion clones were more resistant to stress induced by glucose deprivation. Indeed, MCF7Δ#1 and MCF7Δ#2 clones carrying the engineered deletion were more resistant to apoptosis under glucose deprivation (Figure 5D). To determine which gene mediates this effect, we overexpressed LINC00636 or CD47 individually in parental MCF7 cells using CRISPRa.

Overexpression of CD47, but not LINC00636, partially recapitulated the apoptosis resistance observed in the deletion clones (Figure 5E), indicating that CD47 contributes to the survival phenotype under nutrient stress.

Cellular senescence is another biological process known to confer resistance to cell death under various stresses (27,28). By performing co-expression pattern analyses using RNA-seq data from TCGA available on The Hub for long non-coding RNAs (lncHUB) (29), we predicted that LINC00636 expression is associated with cellular senescence (Table S1). To test this experimentally, we overexpressed LINC00636 by CRISPRa in MCF7, T47D, and HCC1954 cells and assessed senescence using a senescence-associated β-galactosidase assay. LINC00636 overexpression significantly increased the number of senescent cells in MCF7 and T47D, but not in HCC1954 cultures (Figure 5F and S3D). Moreover, in MCF7 cells, LINC00636 overexpression upregulated IL6 and IL8, key components of the Senescence-Associated Secretory Phenotype (SASP) (30), whereas this effect was not observed in T47D cells (Figure 5G and S3E). This difference may be explained by the fact that, unlike MCF7 cells (31), T47D cells express higher levels of the tumor suppressor and cyclin-dependent kinase inhibitor p16, which can induce senescence independently of the SASP (32,33). Together, these findings indicate that LINC00636 promotes a senescent phenotype in breast cancer models, a process that may contribute to the integral spheroid morphology and survival advantage observed in the 3D cultures.

Next, to test whether the insertion variant has an effect on tumor growth, we performed in vivo experiments using parental MCF7 cells or the MCF7Δ#1 and MCF7Δ#2 clones. We injected the breast cancer cells in the mammary fat pad of 5 weeks old female nude mice and monitored tumor growth using a caliper until tumors reached the endpoint size, after which tumors were collected for flow cytometry analysis (Figure S4A). We did not observe significant differences in tumor growth of MCF7 cells carrying the insertion when compared to the engineered clones carrying the deletion under the tested conditions (Figure S4B). As CD47 is known to inhibit macrophage-mediated phagocytosis of overexpressing cells, and nude mice provide a model in which macrophages remain functional despite the absence of T cells, we investigated whether the insertion variant could influence the tumor microenvironment, specifically macrophage behavior. To this end, we analyzed the immune cell composition of the collected tumors by flow cytometry using the gating strategy shown in Figure S4C. We observed a significant decrease in the percentage of CD80⁺ macrophages in tumors derived from the MCF7Δ#1 and MCF7Δ#2 clones compared to those derived from parental MCF7 cells (Figure 5H and S2D). CD80⁺ macrophages are important regulators of the tumor immune microenvironment, as CD80 expression is typically associated with pro-inflammatory macrophages that support anti-tumor activity by producing cytokines that enhance T cell activation (34,35). Thus, our results show that absence of the insertion decreases breast tumor infiltration of pro-inflammatory macrophages.

### Deletion of the insertion reveals a LINC00636-regulated program of senescence and immune modulation

Next, we asked whether deleting the insertion releases an intrinsic downstream program that is controlled by the insertion. To answer this question, we performed RNA-seq experiments in parental MCF7 cells or the clones carrying the engineered deletion. To identify transcriptional changes driven by the insertion, we focused our analysis on the differentially expressed genes (DEGs) that showed the same trend in both MCF7Δ#1 and MCF7Δ#2 clones compared to parental cells: 535 genes were upregulated and 340 genes were downregulated in both MCF7Δ#1 and MCF7Δ#2 compared to control cells (Figure S6A and Table S2). Next, we clustered the DEGs into pathways using the Reactome pathway database as the reference, which curates biological pathways based on known molecular interactions (36). Reactome pathway analyses for the DEGs showed enrichment for genes with roles in extracellular matrix organization and interferon signaling.

Since deleting the insertion leads to overexpression of LINC00636 (Figure 4C), and little is known about its regulatory role in breast cancer, we aimed to investigate the downstream gene program activated by LINC00636 upregulation resulting from the absence of the insertion. To approach this, we first performed RNA-seq to analyze the transcriptomic changes driven by LINC00636 overexpression in MCF7 cells (using the same CRISPRa method as in Figure 2C). We focused our analysis on the DEGs that showed the same trend when LINC00636 was overexpressed using two different sgRNAs, compared to scramble sgRNA-transfected control cells. We identified 354 upregulated genes and 324 downregulated genes in LINC00636-overexpressing cells (Figure S6B and Table S3). Reactome pathway analyses for the DEGs showed an enrichment in genes involved in cell cycle and interferon signaling. Next, we compared this information to the RNA-seq data obtained after deleting the insertion in MCF7 cells, to identify DEGs that are common to both RNA-seq studies (Figure 6A). We found 26 DEGs common to both RNA-seq datasets, indicating that their expression is affected by deleting the insertion and the resulting upregulation of LINC00636 (Figure 6B). Of these 26 DEGs, 18 were upregulated and 8 were downregulated. Many of the upregulated DEGs have known roles in breast cancer biology, such as apoptosis resistance (e.g., ANGPTL4), resistance to therapy (e.g., KRT17, SALL4, SLC7A5), pro-inflammatory SASP (CXCL8, which codes for IL8), and in tumor immunomodulatory signaling, including transcription factors STAT1 and JUN and chemokine CXCL17 (Table S4). Changes in gene expression for many DEGs were confirmed by qPCR in MCF7Δ#1 and MCF7Δ#2 engineered clones and LINC00636-overexpressing MCF7 cells (Figure S6C).

**Figure 6.**
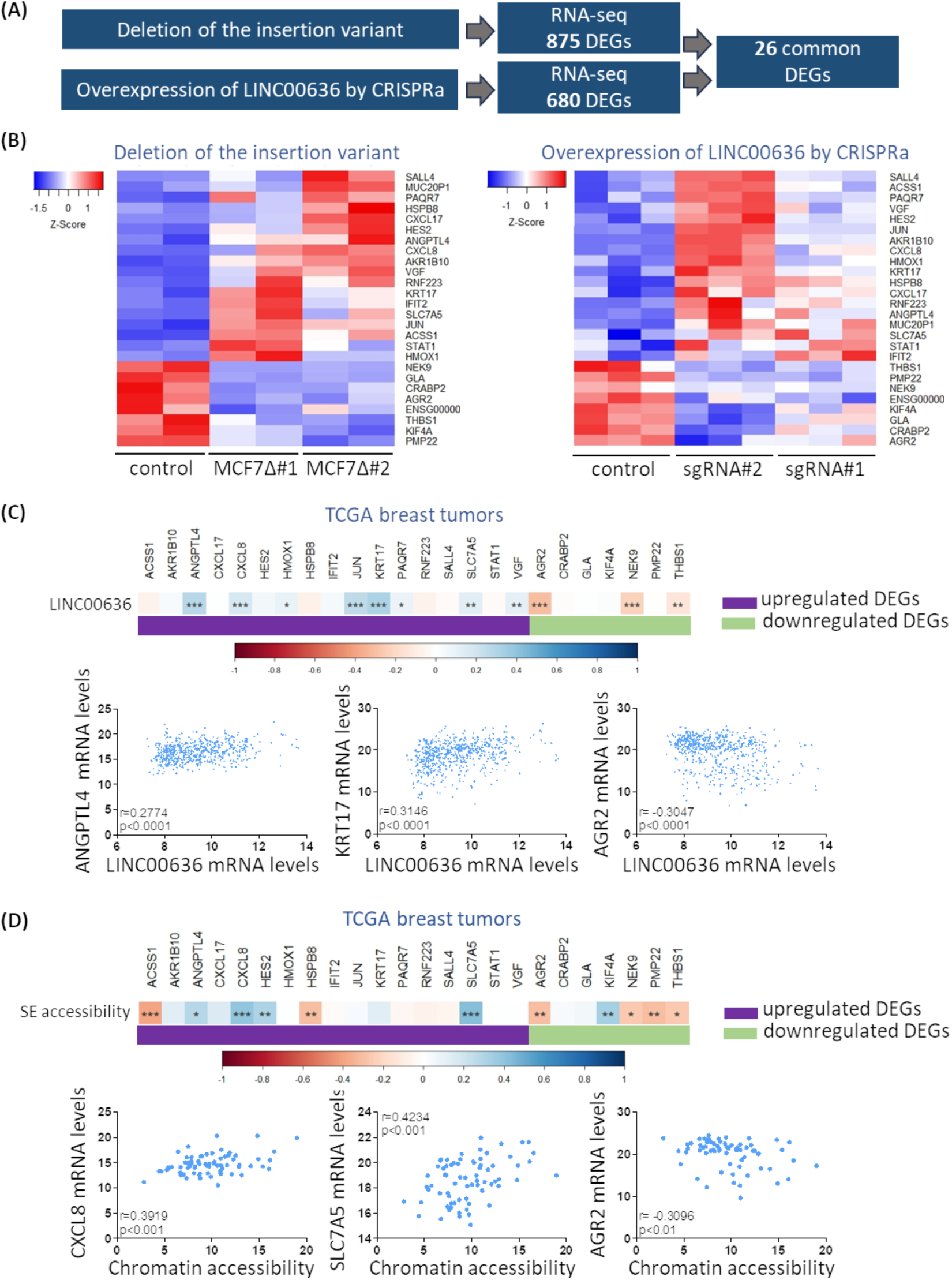
Deletion of the insertion unleashes a downstream transcriptional program through LINC00636 overexpression. (A) Schematic representation of RNA-seq pipeline to identify common DEGs upon deletion of the insertion variant and overexpression of LINC00636 by CRISPRa in MCF7 cells. (B) Heatmaps showing 26 DEGs upon deletion of the insertion variant and overexpression of LINC00636 by CRISPRa in MCF7 cells. (C) Heatmap and representative plots showing LINC00636 RNA levels correlate with RNA expression levels of several DEG in TCGA breast tumors. Violet and green bars indicate, respectively, upregulated and downregulated DEGs found in our RNA-seq analyses. Pearson correlation coefficient, r, is shown in blue/red scale. *p<0.05, **p<0.01, ***p<0.001, n=576. (D) Heatmap and representative plots showing accessibility of the bifunctional SE locus significantly correlates with RNA expression levels of several DEG in breast tumors. Violet and green bars indicate, respectively, upregulated and downregulated DEGs found in our RNA-seq analyses. Pearson correlation coefficient is shown in blue/red scale. *p<0.05, **p<0.01, ***p<0.001, n=72.

We next used TCGA RNA-seq data to investigate whether these regulatory patterns are also observed in breast cancer patients. To approach this, we correlated LINC00636 RNA levels with available RNA levels for 24 DEGs (RNA levels for two DEGs, a pseudogene and an unknown gene, were not available). Our results show that RNA levels of several DEGs are significantly correlated with LINC00636 expression levels, and the direction of the correlation (i.e., positive or negative correlation) matches the one observed in our RNA-seq analyses (Figure 6C and Table S5). Moreover, chromatin accessibility at the bifunctional SE locus in breast tumors significantly correlates with RNA expression of several DEGs (Figure 6D and Table S6).

Our findings show that deletion of the insertion variant within a bifunctional SE increases LINC00636 expression, which in turn activates tumor-intrinsic programs of senescence and pathways that modulate the immune microenvironment (including subsets previously linked to protumorigenic activity), as well as therapy resistance. Thus, our data indicate that the presence of the insertion appears to restrain these pathways, consistent with a less tumorigenic transcriptional state in breast cancer.

## DISCUSSION

Our study demonstrates that a bifunctional SE upregulates LINC00636 and CD47 expression in breast cancer. Moreover, our work shows that a germline InDel variant located on the core element E5 within the SE locus modulates chromatin accessibility (hence, the SE) and the expression of the SE’s target genes. We found that transcription of LINC00636 and, to a lesser extent, of CD47 is regulated by the active SE, as evidenced by: 1) the marked reduction in LINC00636 RNA levels upon BET inhibition in SE-containing breast cancer cell lines, and 2) the significant transcriptional upregulation of LINC00636 after targeting the SE with CRISPRa to test the physical interaction between the SE and TSS of target genes. This suggests a hierarchical interaction between the SE and its target genes, leading to a preferential activation of LINC00636 by the bifunctional SE, potentially influenced by factors such as chromatin looping, and transcription factor occupancy. Such hierarchical interactions have been reported in murine mammary epithelium, where a bifunctional SE primarily upregulates *Wap* expression while modestly regulating the transcription of a secondary target gene, *Ramp3* (37). These findings highlight the role of bifunctional SEs in precisely regulating target genes. Our results also underscore the critical role of core elements within SEs in regulating gene expression in cancer. Through analyses of the TCGA breast cancer patients’ dataset, we identified the core element E5 as the only accessible region within the bifunctional SE significantly correlated with LINC00636 expression in breast tumors, whereas CD47 expression correlates with chromatin accessibility at several accessible regions within the SE. This difference suggests that LINC00636 expression is more dependent on the activity of a single, highly specific regulatory element E5, while CD47 expression may be influenced by a broader range of accessible regions within the SE.

The identification of the InDel variant within the SE locus provides novel insights into the mechanisms behind genetic variation and modulation of the SE function. The InDel variant introduces an additional layer of complexity to gene regulation mediated by the bifunctional SE, through fine-tune modulation of its chromatin accessibility. While deletion of the insertion in MCF7 cells significantly increased chromatin accessibility at the SE locus and transcriptional expression of LINC00636 and CD47, we found that the insertion allele is associated with reduced chromatin accessibility at the SE locus and low levels of LINC00636 and CD47 gene expression. Genetic variation affecting SE-mediated gene regulation has previously been identified in cancer (15,38). However, our work is the first to show that a common InDel variant modulates a bifunctional SE. Our results also suggest that the insertion may recruit repressive factors or otherwise stabilize a closed chromatin configuration. Alternatively, the deletion could bind an activator for the recruitment of additional factors that open the chromatin at the SE site. While the engineered deletions in the MCF7Δ#1 and MCF7Δ#2 clones are not identical, they still produce the same effect on chromatin opening and upregulation of LINC00636 and CD47. Therefore, it is unlikely that an activator can bind to both engineered DNA sequences.

Through transcriptomics analyses, we identified significant transcriptional changes upon deletion of the insertion in MCF7 breast cancer cells. Since deleting the insertion upregulates LINC00636 and, to a lesser extent, CD47, we speculated that the observed transcriptomic changes might be driven mostly by LINC00636 overexpression and not CD47 overexpression. Long intergenic non-coding RNAs have previously been characterized as RNAs that regulate the expression of local and distal target genes through cis- and trans-acting mechanisms (24). We found that LINC00636 overexpression by CRISPRa in MCF7 cells mimics a subset of the gene expression changes observed in MCF7Δ#1 and MCF7Δ#2 clones carrying the engineered deletion, confirming that LINC00636 acts as a downstream effector of the insertion variant.

Specifically, the insertion suppresses tumor-promoting pathways by limiting LINC00636 expression. Among the common DEGs found upregulated upon deleting the insertion and through LINC00636 overexpression, we identified genes such as ANGPTL4 (apoptosis resistance), KRT17, SALL4, and SLC7A5 (resistance to therapy), and chemokines like CXCL8 and CXCL17 (immune modulatory signals), which are well-characterized contributors to tumor progression (39–44). Correlation analyses using TCGA data revealed that the expression of several DEGs is significantly associated with LINC00636 levels in breast cancer patients.

Additionally, chromatin accessibility at the bifunctional SE locus correlates with expression of several DEGs in TCGA breast tumors, reinforcing the mechanistic link between the SE, LINC00636, and downstream gene regulation in a clinical context.

More importantly, we showed that LINC00636 overexpression induces a senescent phenotype in breast cancer cell models and the release of IL6 and IL8 SASP in MCF7 cells. Tumor senescent cells, in addition to being characterized by the upregulation of pro-survival pathways (45), are also known to secrete IL6 and IL8, which have been shown to reshape the tumor microenvironment in multiple ways (46–48). For instance, high IL6/IL8 contributes to chronic inflammation and recruitment of immune suppressive cells such as myeloid-derived suppressor cells that support tumor progression (49–51). Altogether, our data reveals, for the first time, that LINC00636 is associated with a protumoral transcriptional program in breast cancer cells. Our findings are particularly relevant, as the role of LINC00636 in cancer remains poorly understood. Previous studies have only described its overexpression in Her2-positive breast cancers compared to adjacent normal mammary tissue (52) and identified high LINC00636 as a biomarker of unfavorable survival in pancreatic ductal adenocarcinoma patients (53).

Our functional in vitro assays and in vivo experiments illuminated the role of the InDel variant in breast cancer biology. We observed a higher proportion of dead cells in spheroids derived from parental MCF7 cells compared to the clones with the engineered deletion, MCF7Δ#1 and MCF7Δ#2, which maintained spheroid integrity for longer periods. In addition, we found that deleting the insertion increased resistance to apoptosis induced by glucose deprivation, an effect partially mediated by upregulation of CD47. Although an anti-apoptotic and pro-survival role for CD47 has been reported in endometrial carcinoma cells (54), it has not been linked to nutrient deprivation. Thus, the delayed cell death and enhanced resistance to nutrient stress observed in the engineered clones with the deletion may contribute to the survival advantage of these tumors. Our findings suggest that the insertion may contribute to a less aggressive tumor state by promoting earlier cell death. Higher cell death within the tumor microenvironment also contributes to the release of damage-associated molecular patterns (DAMPs) and cytokines by dying cells, which in turn recruit and activate pro-inflammatory macrophages (55). In line with this, our in vivo studies revealed a significant increase in CD80+ pro-inflammatory macrophages in tumors derived from parental cells compared to MCF7Δ#1 and MCF7Δ#2 clones. Since CD80 expression is commonly linked to pro-inflammatory macrophages that enhance anti-tumor immunity by secreting inflammatory cytokines and stimulating T cell activation (34,35), our work shows that the insertion is sufficient to attract CD80+ macrophages, thus reinforcing the tumor microenvironment and potentially enhancing immune surveillance. Since we used a mouse model lacking T cells to enable the engraftment of human cancer cells, our studies only provide an examination of T-cell-independent mechanisms, leaving the exploration of T cell effects for further research. Overall, our in vitro and in vivo results indicate that the insertion variant has a protective role in breast cancer cells by promoting tumor cell death and recruiting pro-inflammatory macrophages to tumor sites.

One limitation of this work is that we experimentally studied the InDel role only in tumor cells. It is possible that the InDel, since it is a common germline variant, can also affect LINC00636 and CD47 expression in other cell populations within the tumor microenvironment. However, the LINC00636 and CD47 bifunctional SE seems to be tumor-type or cell-type specific, as it has not been identified in cancer cell lines derived from other solid tumors or blood cancers (13).

Neither has it been found in immune infiltrates from other solid tumors (data not shown). Single-cell analyses could provide deeper insights into the heterogeneity of SE activity and its impact on gene expression in different cell populations within tumors. We also found that the effects observed upon deletion of the insertion appear to be breast cancer–specific, as they were evident in MCF7 and T47D cell lines, both of which are luminal type. The magnitude of these effects also seemed to depend on the SE status.

In conclusion, our results collectively highlight the intricate interplay between genetic variation, chromatin accessibility, and gene regulation in breast cancer. In particular, we identified a protective insertion variant that decreases accessibility of the bifunctional SE, thus attenuating the expression of the protumorigenic noncoding RNA LINC00636 and the immune-evasion, anti-apoptotic gene CD47. Future studies should focus on elucidating the molecular mechanisms through which the insertion decreases chromatin accessibility at the SE locus.

## METHODS

### Identification of highly accessible DNA regions in access-restricted TCGA ATAC-seq data

Access to access-restricted TCGA ATAC-seq data, deposited on dbGaP phs000178.v11.p8, was approved by the National Institute of Health (NIH). BAM files and indexed files were downloaded from the Genomic Data Commons Data Portal (GDC Data Portal, https://portal.gdc.cancer.gov/). Bigwig files were generated with BPM normalization using Deeptools (56). Peak calling was performed using MACS3 with a q-value=0.00001 (57), and coding regions obtained from https://ftp.ebi.ac.uk/pub/databases/gencode/Gencode_human/release_47/ were subtracted from BED files using BEDtools (58). ROSE with default parameters was used to combine all accessible regions that occurred within 12.5kb of each other into clustered regions, to rank the clusters based on accessibility, and to assign clusters to genes (2). To identify highly accessible regions, the accessibility signal for each cluster was plotted against the clusters’ rank, and the elbow point (first inflection point) was calculated using the package ‘pathviewr’ on RStudio.

Next, we calculated a second elbow point for the lower portion of the curve, using the first inflection point as the new maximum value (17). Clusters whose signal was higher than the second elbow point were considered highly accessible.

In our study, we used the ROSE software only to combine and rank clustered regions based on chromatin accessibility, not to predict SEs from ATAC-seq data. ROSE was originally developed to identify SEs—large clusters of enhancers—using ChIP-seq data for epigenetic marks like H3K27ac, H3K4me1, or transcriptional complexes like Med1 (2). Since ATAC-seq detects all accessible chromatin regions without distinguishing between enhancers, repressors, promoters or insulators, applying ROSE to ATAC-seq data can lead to low true positive and high false positive rates for SE identification. Therefore, in the context of ATAC-seq, ROSE is only suitable for ranking highly accessible chromatin regions to compare across samples.

### Analyses of cancer datasets

RNA-seq and clinical data for 1097 breast cancer primary tumors and ATAC-seq peak signal data for 74 breast cancer primary tumors, all available on The Cancer Genome Atlas (TCGA) dataset (25,59), were downloaded from Xenabrowser (60). Chromatin accessibility of the intergenic regulatory region at chr3:107,985,400-107,994,644 was calculated by combining chromatin accessibility of all individual ATAC-seq peaks located within this DNA region. For correlation analyses of ATAC-seq chromatin accessibility and RNA-seq gene expression, only TCGA samples with gene expression data for LINC00636 (n=45) or CD47 (n=72) were considered.

### In silico analysis of gene-enhancers interactions

Gene-enhancer interactions were visualized on UCSC Genome Browser (61) using the tracks “GeneHancer Regulatory Elements and Gene Interactions”, “GeneCards genes TSS (Double Elite)”, and “Interaction between GeneHancer regulatory elements and genes (Double Elite)” (18).

### InDel genotyping of access-restricted TCGA ATAC-seq data

Access-restricted TCGA ATAC-seq data is deposited on dbGaP phs000178.v11.p8. Using ATAC-seq BAM files as input and GRCh38 as reference genome, we performed a genotype calling with the HaplotypeCaller function of GATK with default parameters, and varDict in single-mode with a minimum allele frequency of 0.01 (62,63). In those cases where both software showed different genotypes for the same sample, we visualized the respective BAM file on the IGV genome viewer and classified the genotype manually. Samples with fewer than 6 reads covering the InDel region were excluded from the analysis.

### Population analysis of InDel frequency

InDel frequencies in general population and across ethnicities were obtained from gnomAD v4.0.0 (64).

### Genomic DNA extraction, PCR, and InDel sequencing of cancer cell lines

Genomic DNA was extracted from breast cancer cell lines using the DNeasy Blood & Tissue Kit (Qiagen), following manufacturer’s instructions. DNA samples were PCR amplified using Phusion High-Fidelity PCR Kit (Thermo Scientific), and the following PCR primers: TGGTCACATGGCTTATTGGG, GGAGAAAACACCCTCCCTGTC. PCR was performed using the following method: an initial incubation at 98°C for 30 seconds, 36 cycles of denaturation at 98°C for 15 seconds, followed by annealing at 63.3°C for 15 seconds, followed by extension at 72°C for 15 seconds; and a final extension at 72°C for 2 minutes. PCR products were isolated by gel electrophoresis followed by DNA extraction using GeneJET Gel Extraction Kit (Fisher), according to manufacturer’s instructions. DNA was sequenced at Genewiz (Azenta Life Sciences), using the same PCR primers.

### Cell culture

MCF7, BT549, HCC1954, BT20 and 293T cell lines were obtained from ATCC. T47D and MDAMB231 were kindly provided by Dr. Mary Helen Barcellos-Hoff and Dr. Denise Munoz, respectively, at UCSF. All breast cancer cell lines except for MDAMB231 were cultured in RPMI medium (Gibco). 293T and MDAMB231 cells were cultured in DMEM medium (Gibco). Media were supplemented with 10% heat inactivated FBS (Atlanta Biologicals), 100 units/mL Penicillin and 100 µg/mL Streptomycin (Gibco). In addition, culture medium for BT549 and T47D cells was supplemented with 0.01 mg/mL insulin (Sigma). Cells were grown at 37°C in a humidified atmosphere at 5% CO_2_.

### Drugs

For BET inhibition, cells were treated with 1 µM JQ1 (Sigma) or 1 µM I-BET151 (GSK1210151A, Selleckchem) dissolved as 10 mM in DMSO; DMSO was used as vehicle control.

### Lentiviral transduction for CRISPRa

Breast cancer cells stably expressing dCas9-VP64 were generated using lentiviral particles. Lentiviral particles were produced by co-transfection of 293T cells with the packaging plasmids psPAX2 and pMD2G, along with lenti-EF1a-dCas9-VP64-Puro plasmid (Addgene #99371) (65) or lenti dCAS-VP64_Blast (Addgene #61425) (66) using TurboFect Transfection Reagent (Life Technologies). Lentiviral particles were harvested 48 hours after transfection in the culture media, concentrated using Lenti-X Concentrator (Takara) following manufacturer’s instructions, and used to infect target cells. After 48 hours of incubation, transduced cells were selected with puromycin or blasticidin.

To generate cells with stable overexpression of sgRNAs for CRISPRa, a second round of transduction was performed in cells expressing dCas9-VP64-blasticidin. Target sgRNA were cloned in the plasmid pSLQ1651-sgRNA(F+E)-sgGal4 (Addgene #100549) (67) by Twist Biosciences. Lentiviral particles were generated, used for transduction and cells were selected with puromycin as described above. pSLQ1651-sgRNA(F+E)-sgGal4 plasmid was used as negative control. sgRNAs target sequences were as follows:

sgRNA scramble: AACGACTAGTTAGGCGTGTA

sgRNA SE: AAAATACGGTTACTGTGATT

sgRNA LINC00636: TCTAGATCTCCCTTTGGTGC or CTAGAGCAAAAGCTGCTTGT

sgRNA CD47: GGGCGCCGCGTCAACAGCAG

### sgRNA transfection for CRISPRa

Synthetic modified sgRNAs (Synthego) for the target region or negative control were transfected at a final concentration of 50nM using TransIT-X2 Dynamic Delivery System (MIR 6004, Mirus) following manufacturer’s instructions. For LINC00636 activation, sgRNAs were designed using CRISPick (68). Negative Control sgRNA (Scrambled sgRNA#1) was from Synthego. sgRNAs target sequences were as follows:

sgRNA SE: AAAATACGGTTACTGTGATT

sgRNA#1: TCTAGATCTCCCTTTGGTGC

sgRNA#2: CTAGAGCAAAAGCTGCTTGT

### RNA extraction, cDNA synthesis and qPCR

Total RNA was extracted using RNeasy Plus kit (Qiagen). cDNA was reversed transcribed using SuperScript III First-Strand Synthesis SuperMix (Invitrogen) and then amplified on the QuantStudio 5 Real Time PCR System (Applied Biosystems). Specific primers designed to amplify the gene of interest were combined with cDNA and SYBR Select Master Mix (Applied Biosystems) following the manufacturer’s instructions. qPCR was carried out using the following method: an initial incubation at 50°C for 2 minutes followed by incubation at 95°C for 2 minutes; 40 cycles at 95°C for 10 seconds followed by incubation at 60°C for 30 seconds; and a final step for melting curve generation. Results were analyzed using the comparative Ct method (69). Values were normalized to β-Actin expression. qPCR primers used in this study are listed on Table S7.

### Deletion of the insertion by CRISPR

Synthetic modified sgRNAs (Synthego) were used for deleting the 8bp insertion within the bifunctional SE. For MCF7 cells, the gRNA sequence was AAAATACGGTTACTGTGATT. For other breast cancer cells, gRNA sequences used were AAAATACGGTTACTGTGATT and AGTGAATAGCAGGAAGTATA. SpCas9 protein and the sgRNA were pre-incubated for 15 minutes at 37°C to obtain ribonucleoprotein complexes. Cells were transfected by electroporation of Cas9-sgRNA ribonucleoprotein using an 4D-Nucleofector™ System (Lonza) according to the manufacturer’s instructions. Briefly, 4×10^5^ cells were mixed with ribonucloprotein complexes in 4D-Nucleofector Solution into a 20 μl Nucleocuvette Strip and programs EN-130, CH125 or FF-150 were used for nucleofection. Following nucleofection, cells were diluted into culture medium and expanded. For MCF7, CRISPR clones were obtained by the clonal picking method, which involves seeding cells in very low density so the cells form colonies, and then manually picking colonies off the plate and transferring them to 96-well plates for clonal expansion. Clones with the desired DNA modification were identified by PCR and DNA sequencing. For other cell lines, pooled cells were used after confirming >99% deletion of the insertion by PCR, DNA sequencing, and analysis of knockout score by EditCo ICE software (https://ice.editco.bio/#/).

### ATAC sequencing

ATAC-seq was performed in MCF7, MCF7Δ#1 and MCF7Δ#2 using the ATAC-Seq Kit (Active Motif, Catalog No. 53150), following manufacturer’s instructions. 1×10^5 fresh cells per sample in duplicates were used. After library preparation, samples were QC using the Bioanalyzer High Sensitivity kit. Libraries were paired-end sequenced in a NovaSeqX PE50, at the Center for Advanced Technology at UCSF. Fastq files were aligned to the hg38 reference genome using Bowtie2 (70), and the resulting SAM files were converted to BAM, sorted and indexed using Samtools (71). BAM files were filtered by proper pairs and indexed using Samtools, inferred duplicates were removed using Picard (http://broadinstitute.github.io/picard), and black-listed regions were removed using Bedtools (58). Bigwig files were generated with BPM normalization using Deeptools (56). Peak calling was performed using MACS3 with a q-value=0.001 (57). The SE or promoter regions were quantified on Bigwig files using Megadepth (72).

### Flow Cytometry

Cells were collected, washed in FACS buffer (2% FBS in PBS) and kept on ice for the rest of the protocol. Cells were stained for 1 hour with APC anti-human CD47 Antibody (clone B6H12, eBiosciences) at a 1:100 dilution in FACS buffer. After washing, cells were stained with Live/dead Fixable Violet Dead (Life Technologies) at a 1:7500 dilution in PBS for 20 minutes. After washing, protein expression was quantified as mean fluorescence intensity (MFI) using a Northern Light cytometer (Cytek). Data was analyzed on SpectroFlo software (Cytek).

### Apoptosis assay

Cells were collected, washed with HBSS and stained with APC or FITC Annexin V (Biolegend, Catalogue #640919 or 640905) in Annexin V binding buffer (Biolegend) for 30 minutes at room temperature. After washing, Annexin V positive events were quantified using a Northern Light cytometer (Cytek). Data was analyzed on SpectroFlo software (Cytek).

### 3D cultures

Insertion-deleted MCF7 clones or control cells were seeded in 12-well plates pre-coated with Phenol-Red Free Matrigel (Corning). The coating was achieved by evenly spreading a thin layer of undiluted Matrigel on each well, followed by incubation at 37°C for 1 hour to allow gelation. After gelation, 50,000 cells were seeded per well in medium supplemented with 5% Matrigel. Spheroid formation was monitored, and images were captured on days 7 or 9 using a Nikon Eclipse TS100 microscope. For dead cell visualization and quantification at day 7, cells were stained at 37°C for 5 minutes with 5µM BD Via-Probe™ Green (BD Biosciences), a nucleic acid stain specific for dead and dying cells, and images were captured using a Nikon Eclipse TS100 microscope. Number of dead cells was quantified using ImageJ and normalized by the number of spheroids in each image.

### Prediction of Gene Ontology Processes for LINC00636

Gene Ontology processes for LINC00636 were predicted using lncHUB (https://maayanlab.cloud/lnchub/), based on RNA-seq co-expression from TCGA (29).

### Senescence-associated β-galactosidase assay

Cellular senescence was measured by β-galactosidase staining using the Cellular Senescence Detection Kit (Cell Biolabs, Catalog # 102964-644), following manufacturer’s instructions. Briefly, after washing and fixing, cells were incubated with staining solution overnight at 37°C without CO2. After washing, images were captured using an Echo Revolve microscope. The percentage of senescent (blue) cells was quantified using ImageJ software.

### In vivo tumor formation

All experimental protocols were approved by the UCSF – Institutional Animal Care & Use Committee (UCSF-IACUC). Five-week-old female inbred nude mice (NU/J, strain 002019, JAX) were randomly divided into 3 groups (n=10 per group) and subcutaneously injected on the mammary fat pad with MCF7, MCF7Δ#1 or MCF7Δ#2 cells (5×10^6^ cells/mouse) diluted 1:1 in Matrigel (Corning). Palpable masses were measured twice a week using a caliper and tumor volume was quantified using the formula V = (π.D.d^2)/6, where D is the largest diameter and d is the smallest diameter of the tumor.

### Tumor collection, processing and immune profiling

Tumors were collected at endpoint, cut into small pieces and frozen in FBS containing 10% DMSO until further processing. For immune profiling, frozen vials were thawed, pieces were rinsed in RPMI media four times and minced with small dissection scissors. Minced tissue was incubated with 0.1mg/ml DNase, 4mg/ml Collagenase IV in 5ml RPMI + 2%FBS per sample at 37°C for 30 minutes with shaking. After digestion, enzymes were neutralized with FACS buffer and samples were filtered through a 100 µm cell strainer and washed twice with FACS buffer. 2×10^6 cells per sample were stained with Zombie Aqua Live/Dead stain at 1:1000 in PBS and treated with 1:100 mouse FcγR blocker for 30 minutes on ice. Next, cells were stained for 30 minutes on ice with a mix of the following antibodies in FACS buffer: Anti-CD45-PE, clone I3/2.3, Biolegend; Anti-F4/80-APC, clone BM8, Fisher; Anti-CD11b-BV650, clone M1/70, Biolegend; Anti-CD11c-Alexa Fluor 700, clone N418, Biolegend; Anti-CD80-FITC, clone 16-10A1, Biolegend; Anti-CD86-APC/Fire 750, clone GL-1, Biolegend; Anti-I-A/I-E (MHCII)-BV785, clone M5/114.15.2, Biolegend; Anti-CD282 (TLR2)- PE/Cy7, clone QA16A01, Biolegend; Anti-CD163-PE-eFluor 610, clone TNKUPJ, eBiosciences; Anti-CD206-eFluor 450, clone MR6F3, Life Technologies. After washes, cells were analyzed in a Northern Lights cytometer (Cytek). Data was analyzed using SpectroFlo software (Cytek) and FlowJo V10.10 (BD).

### RNA sequencing

LINC00636 overexpression was carried out by CRISPRa using two LINC00636 TSS-targeting sgRNAs or negative control previously described, and RNA was extracted 48 hours after transfection. Total RNA was extracted using RNeasy PLUS kit (Qiagen) in duplicates or triplicates, and RNA-seq was performed by LC Sciences. Poly(A) RNA sequencing library was prepared following Illumina’s TruSeq-stranded-mRNA sample preparation protocol. RNA integrity was checked with Agilent Technologies 2100 Bioanalyzer. Poly(A) tail-containing mRNAs were purified using oligo-(dT) magnetic beads with two rounds of purification. After purification, poly(A) RNA was fragmented using divalent cation buffer in elevated temperature. Quality control analysis and quantification of the sequencing library were performed using Agilent Technologies 2100 Bioanalyzer High Sensitivity DNA Chip. Paired-ended sequencing was performed on Illumina’s NovaSeq 6000 sequencing system. For transcripts assembly, Cutadapt (73) was first used to remove the reads that contained adaptor contamination, low quality bases and undetermined bases. Then sequence quality was verified using FastQC (http://www.bioinformatics.babraham.ac.uk/projects/fastqc/). HISAT2 (74) was used to map reads to the genome of ftp://ftp.ensembl.org/pub/release-107/fasta/homo_sapiens/dna/. The mapped reads of each sample were assembled using StringTie (75). Then, all transcriptomes were merged to reconstruct a comprehensive transcriptome using perl scripts and gffcompare. After the final transcriptome was generated, StringTie and ballgown (http://www.bioconductor.org/packages/release/bioc/html/ballgown.html) were used to estimate the expression levels of all transcripts. For differential expression analysis of mRNAs, StringTie was used to perform expression level for mRNAs by calculating FPKM. mRNAs differential expression analysis was performed by R package DESeq2 (76). The mRNAs with the parameter of false discovery rate (FDR) below 0.05 and absolute fold change ≥ 2 were considered differentially expressed mRNAs. Only DEGs with which showed the same trend when comparing the two insertion-deleted MCF7 clones with the parental cells or when comparing the two sgRNAs for LINC00636 overexpression with the scramble sgRNA, were included in the analysis. Pathway analysis for differentially expressed genes was done using Reactome.org (36).

### Statistical Analysis

All experiments were done in triplicates unless otherwise indicated. Results were plotted as mean values +/- standard deviation using GraphPad Prism7 and statistically analyzed using two-tailed unpaired T-test or ANOVA followed by Dunnet’s or Bonferroni’s post-test, as appropriate. P-values less than 0.05 were considered statistically significant.

## RESOURCE AVAILABILITY

### Lead contact

Requests for further information and resources should be directed to and will be fulfilled by the lead contact, Paola Betancur.

### Materials availability

This study did not generate new unique reagents.

### Data and code availability

- RNA-seq and ATAC-seq data have been deposited at GEO DataSets as GSE290710 and GSE290709 and are publicly available as of the date of publication.
- The TCGA cancer patient data sets analyzed in this study are available in the GDC Data Portal repository, https://portal.gdc.cancer.gov/repository. For this study, publicly available ATAC-seq and RNA-seq TCGA data were downloaded from UCSC XenaBrowser Platform at https://xenabrowser.net/.
- Access-restricted TCGA ATAC-seq data analyzed in this study is deposited on dbGaP at https://www.ncbi.nlm.nih.gov/gap/ with accession number phs000178.v11.p8.
- This paper does not report original code.
- Any additional information required to reanalyze the data reported in this paper is available from the lead contact upon request.

## ACKNOWLEDGMENTS

We thank Dr. Mary Helen Barcellos-Hoff and Dr. Denise Muñoz for sharing materials, and Dr. Brian J. Abraham and Dr. Ignacio Schor for advice on data analysis. This research was funded by the NIH National Cancer Institute K22CA226365, California Breast Cancer Research Program, and the Breast Cancer Research Foundation-AACR Career Development Awards to PB.

## AUTHOR CONTRIBUTIONS

Conceptualization, C.D., A.T., and P.B.; methodology, C.D., A.T., D.A., A.T.,A.S., V.O.M., A.R.L., M.E., H.Z, and P.B.; software, C.D., A.S., and V.O.M.; validation, C.D., A.T., D.A., A.T., A.R.L., M.E.; formal analysis, C.D., A.T., D.A., A.T., A.S., and V.O.M; investigation, C.D., A.T., D.A., A.T.,A.S., V.O.M., A.R.L., M.E.; resources, C.P., H.G.V., I.W., and P.B.; writing—original draft preparation, C.D., A.T., and P.B.; writing—review and editing, C.D., A.T., I.W., and P.B.; visualization, C.D., A.T., D.A., A.T. and P.B.; supervision, C.D. and P.B.; project administration, P.B.; funding acquisition, P.B. All authors have read and agreed to the published version of the manuscript.

## SUPPLEMENTAL INFORMATION

Document S1. Figures S1–S5, Tables S1 and S5–S7

Table S2. Excel file with DEGs identified after deleting the insertion in MCF7 cells

Table S3. Excel file with DEGs identified upon LINC00636 overexpression in MCF7 cells

Table S4. Excel file containing information about the known roles of DEGs in breast cancer

## FIGURE LEGENDS

**Figure S1.** LINC00636 and CD47 RNA expression levels decrease upon BET inhibition in breast cancer cell lines. Cells were treated with 1 μM JQ1 or I-BET151 for 6 or 24 hours and mRNA levels were analyzed by qPCR. DMSO was used as vehicle control. Two-way ANOVA was performed, *p<0.05, **p<0.01, ***p<0.001.

**Figure S2.** Effects of deleting the insertion in breast cancer cells, related to Figure 4. (A) Schematic representation of the dual-sgRNAs CRISPR strategy used to delete the insertion in T47D, HCC1954, BT20 and MDAMB231 cells. The insertion sequence is shown in gray, target sequences for sgRNAs A and B are indicated with solid lines above, PAM sequences with a dotted lines, and red triangles mark the Cas9 cut sites, resulting in a 22bp deletion that contains the insertion. (B) Table showing subtype, SE status, InDel genotype across additional breast cancer cells used for CRISPR deletion of the insertion. (C) LINC00636 and CD47 RNA expression in BCCL upon deletion of the insertion. Upon nucleofection of ribonucleoproteins containing Cas9 and sgRNAs A and B or sgRNA control, cells were grown for one week and mRNA levels were analyzed by qPCR. Two-way ANOVA, ***p<0.001.

**Figure S3.** In vitro effects of deleting the insertion in breast cancer cells, related to Figure 5. (A) MCF7Δ#1 and MCF7Δ#2 clones lose spheroid integrity at a later timepoint compared to MCF7 cells in 3D cultures. MCF7Δ#1 and MCF7Δ#2 clones or MCF7 control cells were seeded on plates pre-coated with Matrigel and imaged at day 9. Three representative images per condition are shown. Scale bars represent 100 μm. (B) LINC00636 and CD47 RNA expression in MCF7 with stable CRISPRa of the bifunctional SE. mRNA levels of MCF7 cells expressing dCas9-VP64 and sgRNAs scramble control or targeting the E5 core element within the SE were analyzed by qPCR. Two-way ANOVA, ***p<0.001. (C) CRISPRa of LINC00636 or CD47 in MCF7 cells does not protect spheroid integrity in 3D cultures. MCF7 cells expressing dCas9-VP64 and sgRNAs scramble control or targeting LINC00636 or CD47 TSSs were seeded on plates pre-coated with Matrigel and imaged at day 7. Scale bars represent 100 μm. mRNA levels were measured by qPCR. ANOVA, ***p<0.001. CD47 protein levels were quantified by flow cytometry. Unpaired t-test, p<0.01. (D) LINC00636 overexpression increases senescence in T47D. T47D and HCC1954 cells expressing dCas9-VP64 and sgRNAs scramble control or sgRNAs targeting LINC00636 TSS were studied. 48 hours post-seeding, senescence was analyzed by b-galactosidase staining and microscopy. Left panel shows representative images. Scale bars represent 110 μm. The plot on the right shows LINC00636 mRNA levels in T47D and HCC1954 cells expressing dCas9-VP64 and sgRNAs scramble control or targeting LINC00636 TSSs analyzed by qPCR. Two-way ANOVA, *p<0.05, ***p<0.001. (E) LINC00636 overexpression does not affect expression of senescence markers IL1A, IL1B, IL6 and IL8 in T47D. T47D cells expressing dCas9-VP64 and sgRNAs scramble control or sgRNAs targeting LINC00636 TSS were studied. 48 hours post-seeding, expression of senescence markers was analyzed by qPCR.

**Figure S4.** In vivo effects of deleting the insertion in MCF7 cells, related to Figure 5. (A) Schematic representation of in vivo experiment. Female nude mice were subcutaneously injected on the mammary fat pad with MCF7, MCF7Δ#1 or MCF7Δ#2 cells (n=10 per group). Tumor growth was monitored twice a week using a caliper. At endpoint, tumors were collected and immune profiled by flow cytometry. (B) Tumor growth is not affected by deleting the insertion in MCF7 cells. 5-week-old female nude mice were subcutaneously injected on the mammary fat pad with MCF7, MCF7Δ#1 or MCF7Δ#2 cells (n=10 per group) and tumor size was measured twice a week. (C) Gating strategy for immune profiling of MCF7 tumors. (D) Infiltration of CD45+ cells, macrophages, dendritic cells (DCs), and CD86+, MHCII+, TLR2+, CD206+ or CD163+ macrophages is similar in tumors derived from insertion-deleted MCF7Δ#1 and MCF7Δ#2 clones compared to parental MCF7 cells.

**Figure S5.** Common DEGs upon deletion of the insertion variant and overexpression of LINC00636 by CRISPRa in MCF7 cells. (A) Pie chart showing DEGs from RNA-seq analysis upon deletion of the insertion variant in MCF7 cells. Enrichment of genes involved in Extracellular matrix organization and Interferon signaling was observed through Reactome Pathways Analysis of DEGs. n=2. (B) Pie chart showing DEGs from RNA-seq analysis upon overexpression of LINC00636 by CRISPRa in MCF7 cells. Enrichment of genes involved in Cell cycle and Interferon signaling was observed through Reactome Pathways Analysis of DEGs. n=3. (C) qPCR analyses validate gene expression changes identified through RNA-seq upon deletion of the insertion variant and overexpression of LINC00636 in MCF7 cells. ANOVA was performed, *p<0.05, **p<0.01, ***p<0.001.

