## supplemental figures and tables for "An InDel Genomic Variant within a Bifunctional Super-Enhancer for LINC00636 and CD47 Regulation in Breast Cancer"

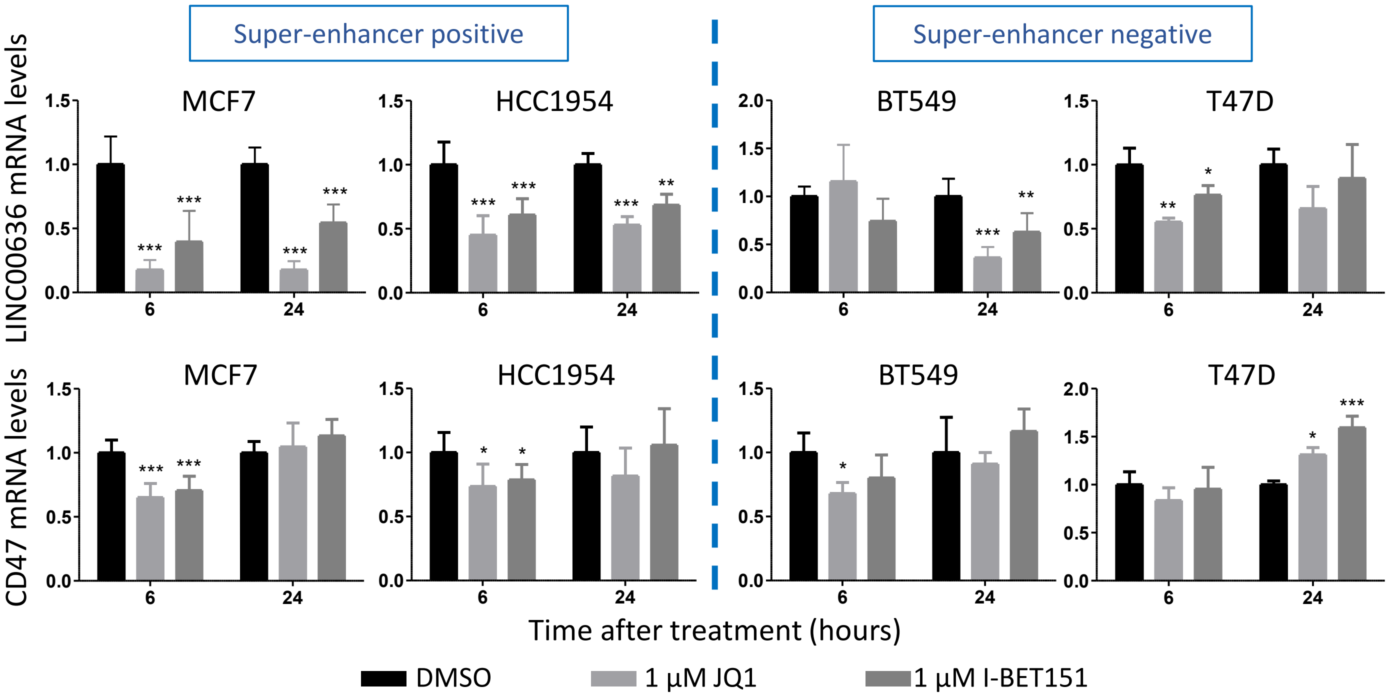


Figure S1. LINC00636 and CD47 RNA expression levels decrease upon BET inhibition in breast cancer cell lines, related to Figure 2. Cells were treated with 1 µM JQ1 or I-BET151 for 6 or 24 hours and mRNA levels were analyzed by qPCR. DMSO was used as vehicle control. Two-way ANOVA, *$p<0.05$, **$p<0.01$, ***$p<0.001$.

**(A)**

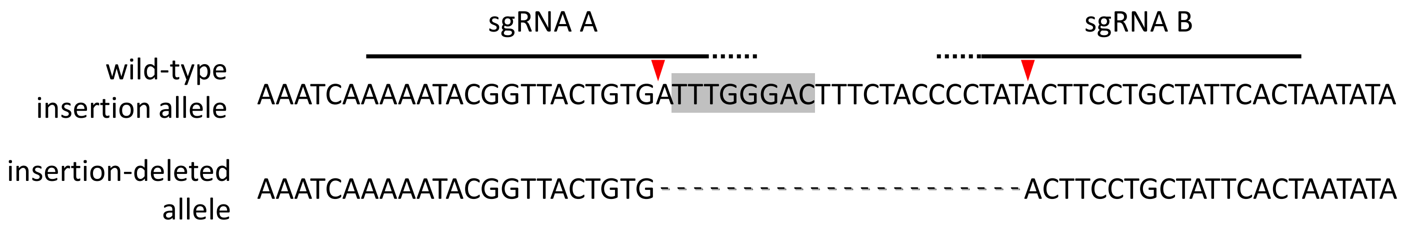


**(B)**

| Cell line | Breast cancer subtype | SE status | InDel genotype |
|---|---|---|---|
| T47D | Luminal | - | Insertion |
| HCC1954 | Her2 | + | Heterozygous |
| BT20 | Basal | - | Insertion |
| MDAMB231 | Basal | - | Insertion |

**(C)**

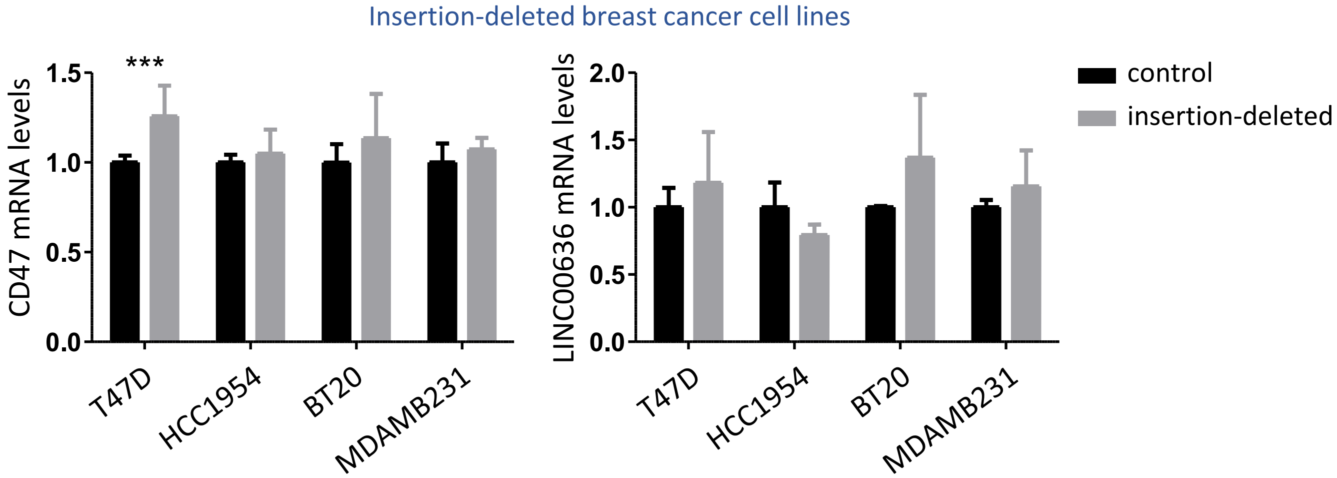


Figure S2. Effects of deleting the insertion in breast cancer cells, related to Figure 4. (A) Schematic representation of the dual-sgRNAs CRISPR strategy used to delete the insertion in T47D, HCC1954, BT20 and MDAMB231 cells. The insertion sequence is shown in gray, target sequences for sgRNAs A and B are indicated with solid lines above, PAM sequences with a dotted lines, and red triangles mark the Cas9 cut sites, resulting in a 22bp deletion that contains the insertion. (B) Table showing subtype, SE status, InDel genotype across additional breast cancer cells used for CRISPR deletion of the insertion. (C) LINC00636 and CD47 RNA expression in BCCL upon deletion of the insertion. Upon nucleofection of ribonucleoproteins containing Cas9 and sgRNAs A and B or sgRNA control, cells were grown for one week and mRNA levels were analyzed by qPCR. Two-way ANOVA, ***p<0.001.

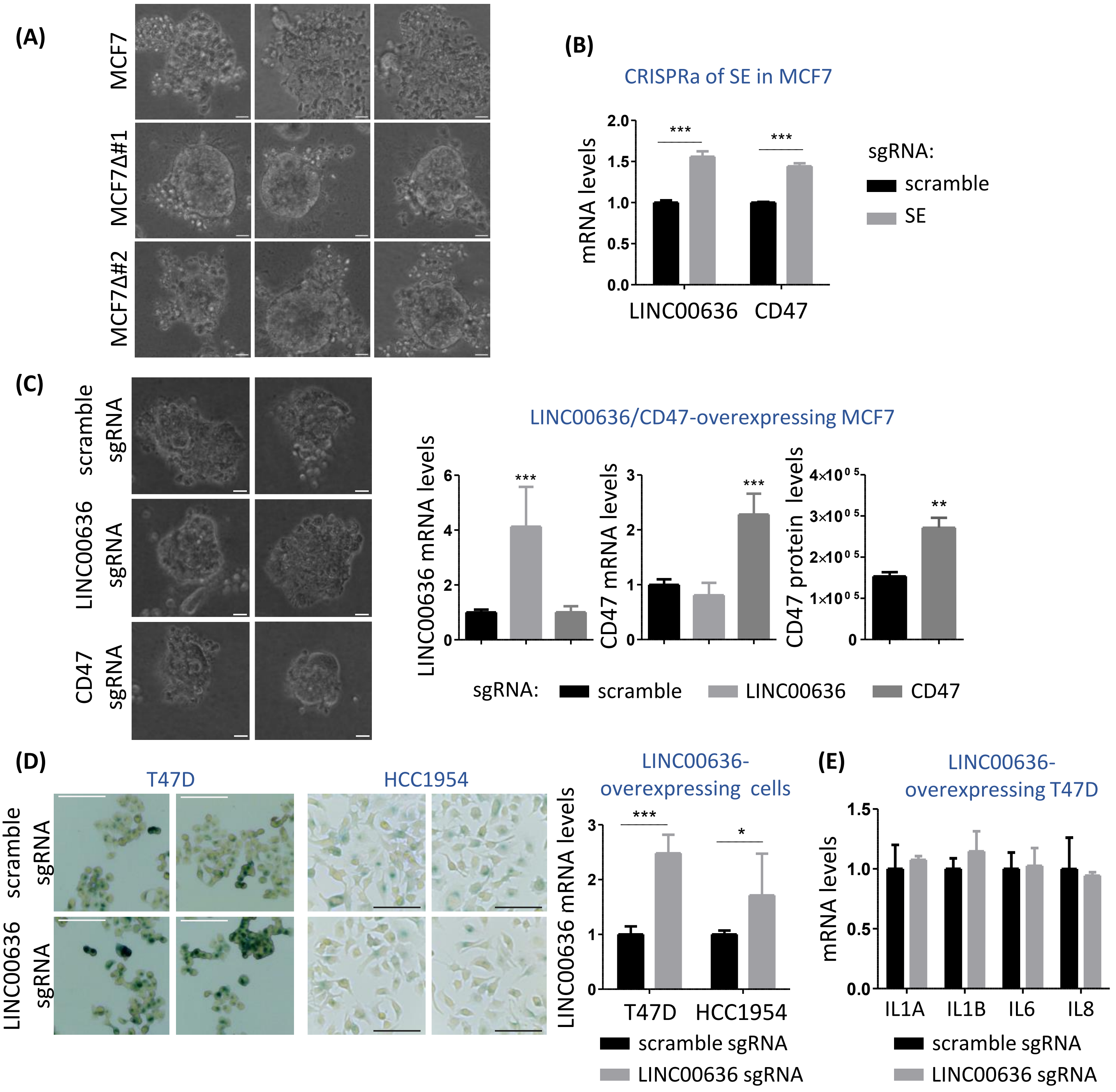


Figure S3. In vitro effects of deleting the insertion in breast cancer cells, related to Figure 5. (A) MCF7Δ#1 and MCF7Δ#2 clones lose spheroid integrity at a later timepoint compared to MCF7 cells in 3D cultures. MCF7Δ#1 and MCF7Δ#2 clones or MCF7 control cells were seeded on plates pre-coated with Matrigel and imaged at day 9. Three representative images per condition are shown. Scale bars represent 100 µm. (B) LINC00636 and CD47 RNA expression in MCF7 with stable CRISPRa of the bifunctional SE. mRNA levels of MCF7 cells expressing dCas9-VP64 and sgRNAs scramble control or targeting the E5 core element within the SE were analyzed by qPCR. Two-way ANOVA, ***p<0.001. (C) CRISPRa of LINC00636 or CD47 in MCF7 cells does not protect spheroid integrity in 3D cultures. MCF7 cells expressing dCas9-VP64 and sgRNAs scramble control or targeting LINC00636 or CD47 TSSs were seeded on plates pre-coated with Matrigel and imaged at day 7. Scale bars represent 100 µm. mRNA levels were measured by qPCR. ANOVA, ***p<0.001. CD47 protein levels were quantified by flow cytometry. Unpaired t-test, p<0.01. (D) LINC00636 overexpression increases senescence in T47D. T47D and HCC1954 cells expressing dCas9-VP64 and sgRNAs scramble control or sgRNAs targeting LINC00636 TSS were studied. 48 hours post-seeding, senescence was analyzed by b-galactosidase staining and microscopy. Left panel shows representative images. Scale bars represent 110 µm. The plot on the right shows LINC00636 mRNA levels in T47D and HCC1954 cells expressing dCas9-VP64 and sgRNAs scramble control or targeting LINC00636 TSSs analyzed by qPCR. Two-way ANOVA, *p<0.05, ***p<0.001. (E) LINC00636 overexpression does not affect expression of senescence markers IL1A, IL1B, IL6 and IL8 in T47D. T47D cells expressing dCas9-VP64 and sgRNAs scramble control or sgRNAs targeting LINC00636 TSS were studied. 48 hours post-seeding, expression of senescence markers was analyzed by qPCR.

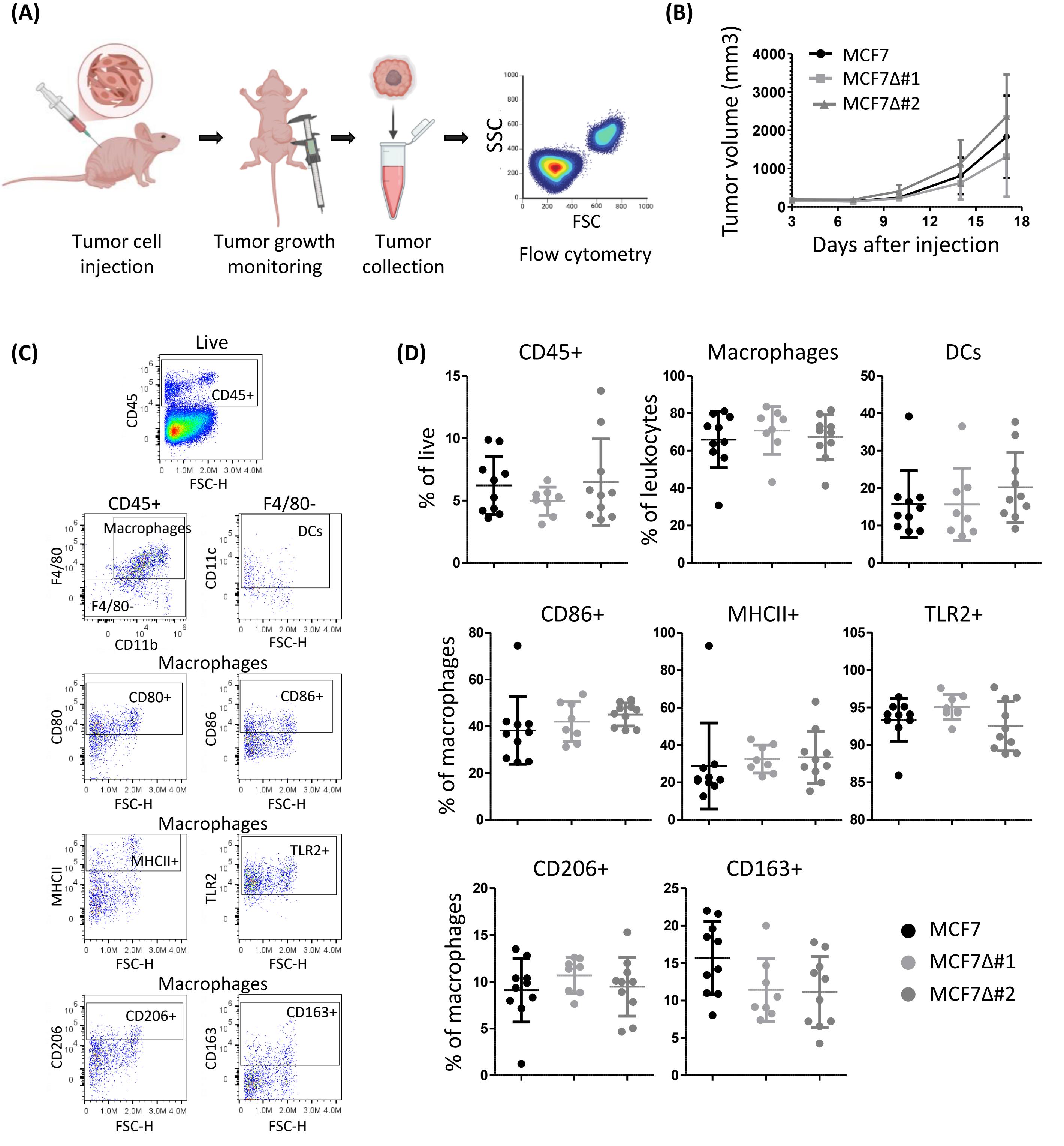


Figure S4. In vivo effects of deleting the insertion in MCF7 cells, related to Figure 5. (A) Schematic representation of in vivo experiment. Female nude mice were subcutaneously injected on the mammary fat pad with MCF7, MCF7Δ#1 or MCF7Δ#2 cells (n=10 per group). Tumor growth was monitored twice a week using a caliper. At endpoint, tumors were collected and immune profiled by flow cytometry. (B) Tumor growth is not affected by deleting the insertion in MCF7 cells. 5-week-old female nude mice were subcutaneously injected on the mammary fat pad with MCF7, MCF7Δ#1 or MCF7Δ#2 cells (n=10 per group) and tumor size was measured twice a week. (C) Gating strategy for immune profiling of MCF7 tumors. (D) Infiltration of CD45+ cells, macrophages, dendritic cells (DCs), and CD86+, MHCII+, TLR2+, CD206+ or CD163+ macrophages is similar in tumors derived from insertion-deleted MCF7Δ#1 and MCF7Δ#2 clones compared to parental MCF7 cells.

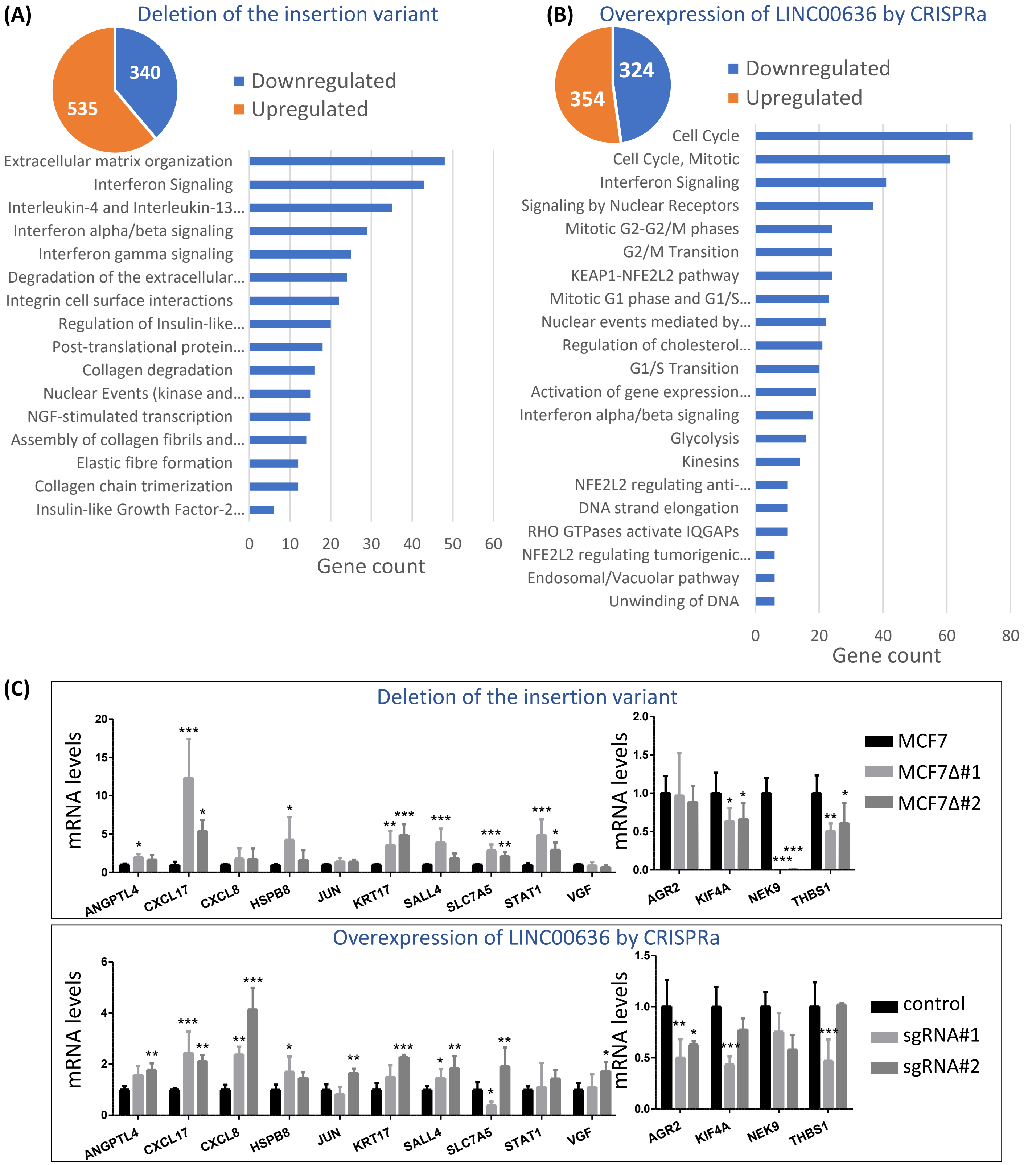


Figure S5. Common DEGs upon deletion of the insertion variant and overexpression of LINC00636 by CRISPRa in MCF7 cells, related to Figure 6. (A) Pie chart showing DEGs from RNA-seq analysis upon deletion of the insertion variant in MCF7 cells. Enrichment of genes involved in Extracellular matrix organization and Interferon signaling was observed through Reactome Pathways Analysis of DEGs, n=2. (B) Pie chart showing DEGs from RNA-seq analysis upon overexpression of LINC00636 by CRISPRa in MCF7 cells. Enrichment of genes involved in Cell cycle and Interferon signaling was observed through Reactome Pathways Analysis of DEGs, n=3. (C) qPCR analyses validate gene expression changes identified through RNA-seq upon deletion of the insertion variant and overexpression of LINC00636 in MCF7 cells. ANOVA, *p<0.05, **p<0.01, ***p<0.001.

| Gene Ontology Biological Processes Predictions | | |
|---|---|---|
| Rank | Process | z-score |
| 1 | regulation of male gonad development (GO:2000018) | 2.719 |
| 2 | positive regulation of metanephros development (GO:0072216) | 2.675 |
| 3 | positive regulation of cellular senescence (GO:2000774) | 2.532 |
| 4 | positive regulation of cell aging (GO:0090343) | 2.337 |
| 5 | response to gonadotropin (GO:0034698) | 2.249 |
| 6 | prostanoid biosynthetic process (GO:0046457) | 2.142 |
| 7 | regulation of organ morphogenesis (GO:2000027) | 2.131 |
| 8 | tube closure (GO:0060606) | 1.973 |
| 9 | cellular response to gonadotropin stimulus (GO:0071371) | 1.972 |
| 10 | folic acid transport (GO:0015884) | 1.97 |

Table S1. Predicted Gene Ontology Biological Processes of LINC00636, related to Figure 5. Predictions are based on co-expression patterns using gene expression data from 11,284 TCGA RNA-seq samples available from lncHUB (https://maayanlab.cloud/lnchub/). Top 10 predicted processes are shown.

| Gene | Correlation between gene RNA expression and LINC00636 RNA expression (TCGA BRCA, n=576) | | |
|---|---|---|---|
| | Pearson r | P value | P value summary |
| ACSS1 | -0.06867 | 0.0997 | ns |
| AKR1B10 | 0.03815 | 0.3608 | ns |
| ANGPTL4 | 0.2774 | < 0.0001 | *** |
| CXCL17 | 0.006252 | 0.881 | ns |
| CXCL8 | 0.1762 | < 0.0001 | *** |
| HES2 | 0.05389 | 0.1966 | ns |
| HMOX1 | 0.09243 | 0.0265 | * |
| HSPB8 | -0.07497 | 0.0722 | ns |
| IFIT2 | 0.04389 | 0.293 | ns |
| JUN | 0.2574 | < 0.0001 | *** |
| KRT17 | 0.3146 | < 0.0001 | *** |
| PAQR7 | 0.0887 | 0.0333 | * |
| RNF223 | -0.06091 | 0.1443 | ns |
| SALL4 | -0.02495 | 0.5502 | ns |
| SLC7A5 | 0.1282 | 0.0021 | ** |
| STAT1 | -0.02062 | 0.6215 | ns |
| VGF | 0.1168 | 0.005 | ** |
| AGR2 | -0.3047 | < 0.0001 | *** |
| CRABP2 | -0.01049 | 0.8017 | ns |
| GLA | 0.003752 | 0.9284 | ns |
| KIF4A | 0.01332 | 0.7498 | ns |
| NEK9 | -0.2204 | < 0.0001 | *** |
| PMP22 | -0.001663 | 0.9682 | ns |
| THBS1 | -0.1357 | 0.0011 | ** |

Table S5. Correlations between RNA expression of DEGs and LINC00636 in TCGA breast tumors, related to Figure 6. RNA expression levels of several DEG significantly correlate with LINC00636 RNA levels in breast tumors. Pearson correlation coefficient r was calculated. *p<0.05, **p<0.01, ***p<0.001, n=576.

| Gene | Correlation between gene RNA expression and bifunctional SE locus accessibility (TCGA BRCA, n=72) | | |
|---|---|---|---|
| | Pearson r | P value | P value summary |
| ACSS1 | -0.3949 | 0.0006 | *** |
| AKR1B10 | 0.1207 | 0.3124 | ns |
| ANGPTL4 | 0.2862 | 0.0148 | * |
| CXCL17 | 0.1506 | 0.2066 | ns |
| CXCL8 | 0.3919 | 0.0007 | *** |
| HES2 | 0.3046 | 0.0093 | ** |
| HMOX1 | 0.004161 | 0.9723 | ns |
| HSPB8 | -0.304 | 0.0094 | ** |
| IFIT2 | -0.02723 | 0.8204 | ns |
| JUN | 0.03668 | 0.7597 | ns |
| KRT17 | 0.1295 | 0.2784 | ns |
| PAQR7 | -0.04021 | 0.7374 | ns |
| RNF223 | -0.06107 | 0.6103 | ns |
| SALL4 | -0.05149 | 0.6676 | ns |
| SLC7A5 | 0.4234 | 0.0002 | *** |
| STAT1 | 0.007964 | 0.9471 | ns |
| VGF | 0.006135 | 0.9592 | ns |
| AGR2 | -0.3096 | 0.0081 | ** |
| CRABP2 | 0.03916 | 0.744 | ns |
| GLA | 0.05984 | 0.6175 | ns |
| KIF4A | 0.3341 | 0.0041 | ** |
| NEK9 | -0.2609 | 0.0268 | * |
| PMP22 | -0.3089 | 0.0083 | ** |
| THBS1 | -0.2652 | 0.0244 | * |

Table S6. Correlations between RNA expression of DEGs and accessibility of the bifunctional SE locus in TCGA breast tumors, related to Figure 6. RNA expression levels of several DEG significantly correlate with accessibility of the bifunctional SE locus in breast tumors. Pearson correlation coefficient r was calculated. *p<0.05, **p<0.01, ***p<0.001, n=72.

| Gene | Forward (5'-3') | Reverse (5'-3') |
|---|---|---|
| β-Actin | TCCCTGGAGAAGAGCTACG | GTAGTTTCGTGGATGCCACA |
| CD47 | CATGGCCCTCTTCTGATTTC | GGAGGTTGTATAGTCTTCTGATTGG |
| LINC00636 | GTCTCAAAGCACGGATGGGA | AAATGCCGAGATGTCCCAGG |
| ANGPTL4 | CAGGGGTCTCCGCCATTTTT | CGGTTGAAGTCCACTGAGCC |
| CXCL17 | TTCATGACAGTGTCTGGGCTG | CTTTCTGTGGTGCCTTTGGTG |
| CXCL8 | AGTTTTTGAAGAGGGCTGAGA | TGCTTGAAGTTTCACTGGCATC |
| HSPB8 | AGCTTCAAGCCAGAGGAGTTG | ATGCCACCTTCTTGCTGTTTC |
| JUN | ACGGCGGTAAAGACCAGAAG | CCAAGTTCAACAACCGGTGC |
| KRT17 | GCCCGTGACTACAGCCAGT | GGATGTTGGCATTGTCCACG |
| SALL4 | TTGTGGGCGAGCTTTTACCA | TCGATGGCCAACTTCCTTCC |
| SLC7A5 | GTCTGGTGGAAAAACAAGCCC | CCTGCATGAGCTTCTGACAC |
| STAT1 | AGGTCTCAATGTGGACCAGC | ACAAAACCTCGTCCACGGAA |
| VGF | TGACACCAGCTGTCTCCG | CAGCAGAAGGCAGAAGAGGG |
| AGR2 | CATCACTTGGATGAGTGCCCA | AGGACAAACTGCTCTGCCAAT |
| KIF4A | AACTGCAGCCCATTCAGTACC | TGGCATCCTTCTTTGCTGTCT |
| NEK9 | GAGCAGGAGGAACTGCACTA | CCGGGTCAAATCGACTTCCT |
| THBS1 | CTGCTCCAATGCCACAGTTC | GCCACAGCTCGTAGAACAGG |
| IL1A | AGATGCCTGAGATACCCAAAACC | CCAAGCACACCCAGTAGTCT |
| IL1B | TTCGACACATGGGATAACGAGG | CTGGCCGTGGCTCTCTTGG |
| IL6 | AGACAGCCACTCACCTCTTCAG | TTCTGCCAGTGCCTCTTTGCTG |
| IL8 | CTGGCCGTGGCTCTCTTGG | CTTGGCAAAACTGCACCTTCA |

Table S7. Primers used for qPCR, related to Methods.
